# Interannual changes in nutrient and phytoplankton dynamics in the Eastern Mediterranean Sea (EMS) predict the consequences of climate change; results from the Sdot-Yam Time-series station 2018-2022

**DOI:** 10.1101/2024.06.24.600321

**Authors:** Tal Ben-Ezra, Alon Blachinsky, Shiran Gozali, Anat Tsemel, Yotam Fadida, Dan Tchernov, Yoav Lehahn, Tatiana Margo Tsagaraki, Ilana Berman-Frank, Michael Krom

## Abstract

Global climate change is predicted to reduce nutrient fluxes into the photic zone, particularly in tropical and subtropical ocean gyres, while the occasional major storms will result in increased nutrient pulses. In this study the nutrient and phytoplankton dynamics have been determined at a new time-series station in the southeastern Levantine basin of the Eastern Mediterranean Sea (EMS) over 4.5 years (2017-2022). In 2018 and 2019, there was a moderate concentration of residual nitrate and nitrite (N+N) in the photic zone (280-410nM) in winter, resulting in phytoplankton dynamics dominated by cyanobacteria with relatively few picoeukaryotes (280± 90 μgC m^−2^). Winter storm driven mixing was much reduced in 2020 and particularly in 2021, resulting in a lower concentration of N+N in the photic zone, which decreased during summer stratification, such that by August 2021, the N+N was highly depleted (<60 nM) resulting in an integrated phytoplankton biomass of 23 μgC m^−2^. A major storm in December 2021 (Storm Carmel) injected high N+N (750 nM; max = 1090 nM) in the upper 100 m, which stimulated pico and nanophytoplankton biomass (∼2400 μgC m^−2^) and probably increased eukaryotes (diatoms). The pattern of measured silica reinforced our conclusion that we sampled 3 different nutrient and ecosystem states. Phosphate was always at or close to LoD because of rapid uptake by cyanobacteria into their periplasm. These results predict that climate change in the EMS will result in periods of nutrient and phytoplankton depletion (Famine) interrupted by short periods of Mesotrophy (Feast) caused by major storms.

**Highlights:**

– Nutrient dynamics from 4 years of Time-series station in the S.E. Levantine basin
– Defined ecosystem status of normal, depleted and temporarily mesotrophic which are predicted status’ caused by climate change
– Winters with low deep mixing resulted in severely nutrient depleted conditions subsequently
– Major storm and relatively shallow nutricline resulted in temporary mesotrophic status

## 1. Introduction

Global climate change is having important effects on the surface waters of the global ocean and consequently on the entire marine ecosystem (Winder and Sommer 2012). Systematic increases in the temperature and pH of the photic zone are widely observed, although the magnitude of such changes varies locally and regionally (Hurd et al. 2018). Moreover, nutrient fluxes into the surface waters are predicted to change. Increased surface temperatures, particularly in the wide-spread tropical and subtropical gyres, will result in less intense winter mixing and longer periods of seasonal stratification (Behrenfeld et al. 2006). Subsequently, carbon uptake, primary productivity, and the efficiency of the carbon pump (i.e. carbon exported to depth) are expected to decline (Marinov et al. 2010). While longer periods of ocean stability are predicted, a larger number of more intense storms is also forecast in the future oceans (Korty 2022). Depending on the timing of such storms, they are likely to produce major pulses of nutrients into the surface waters and higher carbon uptake and phytoplankton biomass (Shan et al. 2023).

The Eastern Mediterranean sea (EMS) in general and the southeastern Levantine basin in particular, represent an important natural laboratory for globally important biogeochemical processes (Thingstad et al. 2005; Moutin et al. 2012; Krom et al. 2014a; Powley et al. 2017). The southeastern Levantine basin is ultraoligotrophic as a result of its unusual anti-estuarine circulation (Krom et al. 2014a). Surface waters flow in through the straits of Sicily and become progressively more saline as they flow towards the east and down-well in winter to form Levantine intermediate water (LIW), which flows out of the basin. The residence time of this LIW is less than eight years (Roether et al. 1998). LIW flows out carrying inorganic and organic nutrients from the basin, causing it to have very low primary productivity and phytoplankton biomass (Moutin and Raimbault 2002; Powley et al. 2014).

The EMS is also phosphorus (P) depleted (Krom et al. 1991). The nitrate to dissolved inorganic phosphate (N+N:DIP) ratio in deep water is ∼28:1 and the PON:POP and DON:DOP ratios in surface and intermediate waters are also greatly over 16:1 (Krom et al. 2005b; Pujo-Pay et al. 2011). Recent high-sensitivity measurements of N+N:DIP ratios in the photic zone throughout the year confirm that the ratios are often >>16:1 (Ben Ezra et al. 2021). The depletion of P in the basin is primarily attributed to a high N+N:DIP ratio in major external inputs (atmospheric and riverine inputs which have an N:P ratio of >> 16:1; Krom et al., 2010) and low rates of denitrification (Powley et al., 2017). In many ways the EMS acts like an ocean gyre such as the N. Atlantic gyre which is also both P depleted (Wu et al. 2000; Reynolds et al. 2014) and net heterotrophic.

The EMS is particularly vulnerable to climate change (Schroeder et al. 2017). Ozer et al. (2017) calculated an increase in temperature of the upper layers of 0.12 °C year^−1^ over the last decade, while a more recent estimate is somewhat lower 0.05 °C year^−1^ but still substantial (Ozer et al. 2022). This compares with a global average increase of 0.013 °C year^−1^ since 1980 (EPA 2021). Such predicted changes in temperature are expected to have inhibitory effects on native EMS marine species e.g. macroalgae and coccolithophores (D’Amario et al. 2020; Mulas et al. 2022). The effect of climate change such as increased CO_2_ and temperature, on phytoplankton are predicted to be complex with increased carbon uptake rate but conversion to DOC due to the reduced supply of nutrients (Vichi et al. 2003).

To study the effects of climate change on nutrient and phytoplankton dynamics in the EMS, we have set up a time-series station at a location with a water depth of ∼800 m off the Israeli coast. The aim of this station is to develop a series of physical, chemical, and biological measurements to identify long-term changes, particularly examining high sensitivity dissolved nutrient concentrations together with partial phytoplankton species and biomass. At present we visit the station monthly-quarterly, and a series of physical, chemical and a limited number of biological measurements are made on each cruise visit. Analysis of the initial 4.5 years of data from this previously undocumented pelagic station reveals the presence of characteristic seasonal and interannual variations. These findings provide a baseline for understanding the dynamics of this region and suggest potential influences of climatic forces. We proceed to identify three distinct states that this system may adopt, offering insights into the future trajectory of the EMS marine environment. Furthermore, the identified dynamics and state transitions that may hold broader implications for the understanding and prediction of change in other marine systems facing similar environmental pressures.

## 2. Sampling and methods

### 2.1. Sampling

This work combines samples we collected from January 2018 to January 2019 at THEMO2 observatory, located at 32.79 N 34.38 E which is ∼27 km offshore at a depth of 1450 m, as detailed in Ben-Ezra et al; 2021 and Reich et al; 2022, and at a pelagic station (N800) located at a depth of 832 m ∼15 km offshore the Israeli coast starting January 2020, which has become the long-term time-series station continuing to the present. N800 water samples were collected monthly or every 2-3 months. The sampling times and parameters are detailed in Table S1. The N800 pelagic station, located at 32.52 N 34.72 E shown in Figure 1, was chosen because it was located in deep water and is known to have insignificant effects of lateral nutrient inputs from the coastal shelf. As observed through remote sensing, filaments have transported water from the coast only in July 2018 and 2020. However, this has minor effects on the time-series station since coastal and offshore waters exhibit very similar biogeochemistry (Ben-Ezra et al., 2023).

**Figure 1:**
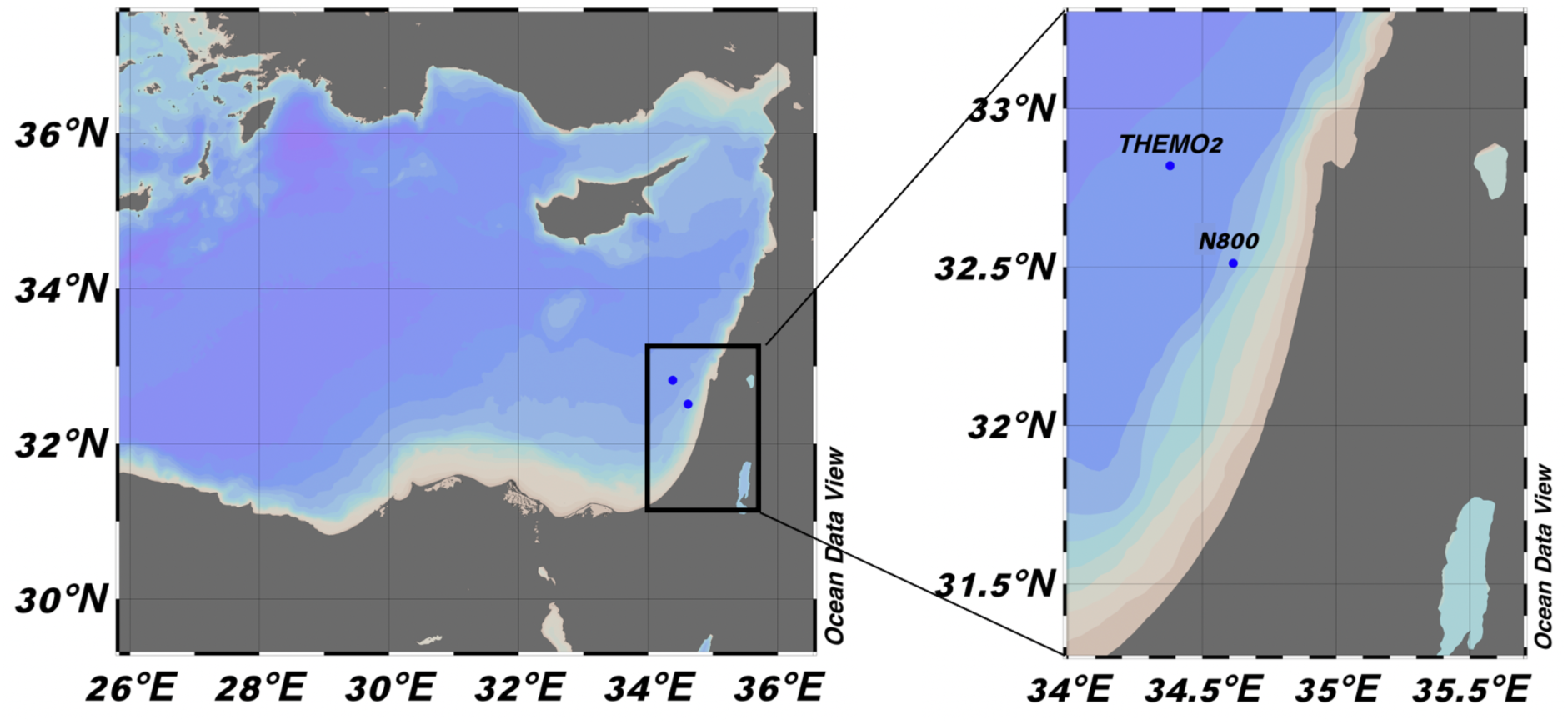
Sampling locations included in this study. Station THEMO2 located at 32.79 N 34.38 E and station N800 located at 32.52 N 34.72 E.

**Figure 2:**
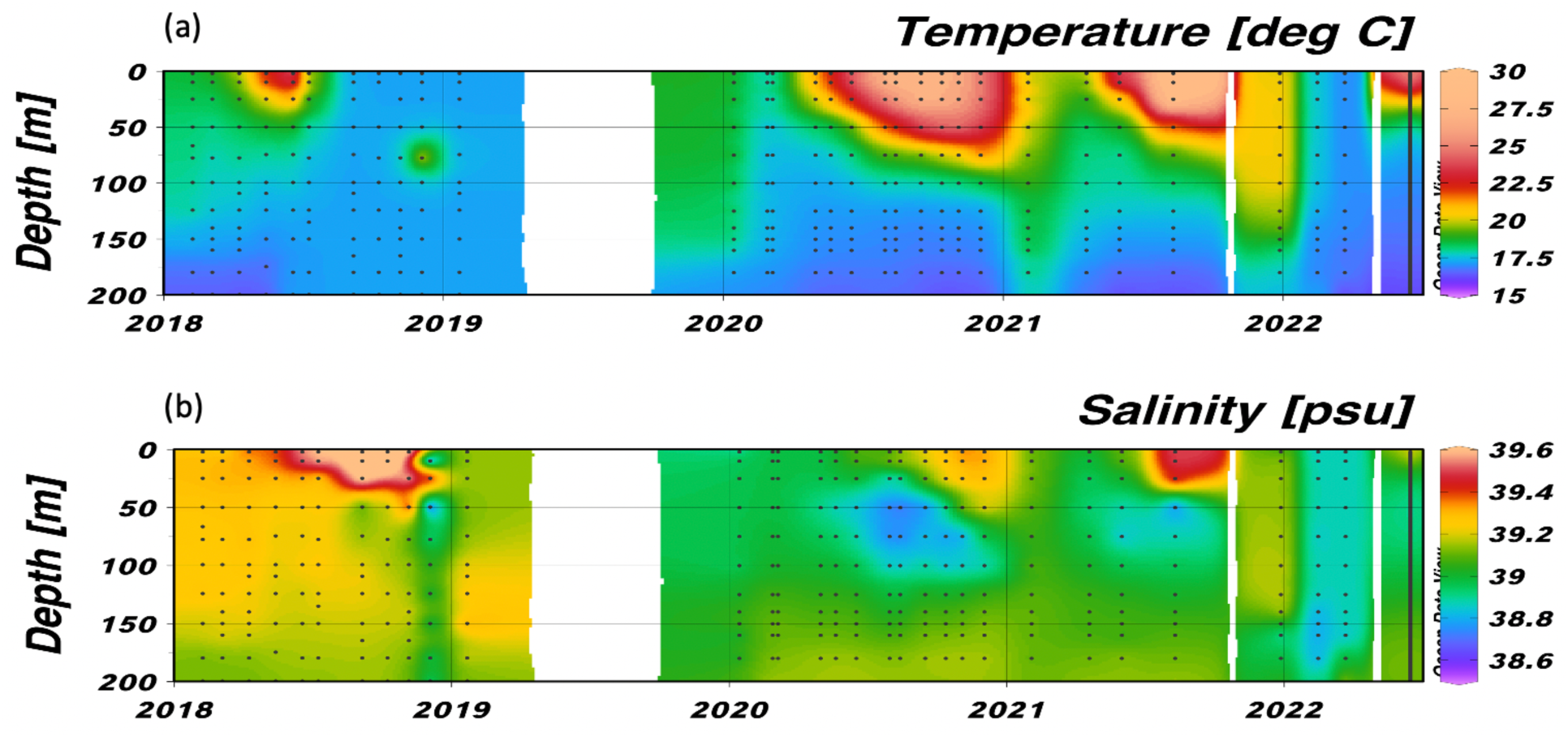
Temperature (a) and salinity (b) time-series at the two EMS pelagic stations. Data shown is from the top 200 m. The data from 2018-2019 are from THEMO2 station (Ben-Ezra et al., 2021) while from 2020 onwards the samplings were from station N800. The sampling depths are marked as dots, contouring was obtained using ODV weighted average gridding set for 25 permille x scale-length and 30 permille y scale-length.

**Figure 3:**
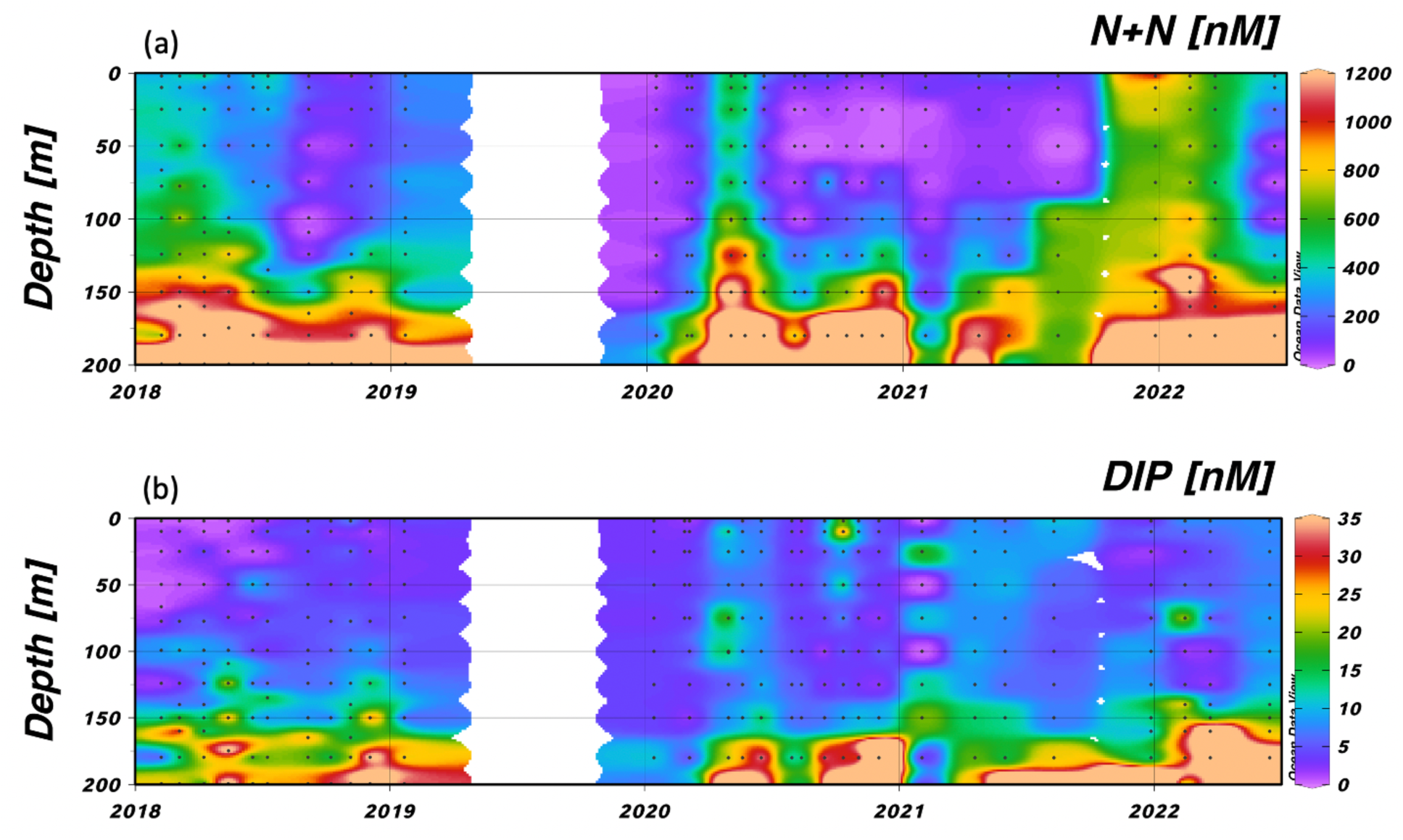
(a) Nitrate+nitrite (N+N) and (b) dissolved inorganic phosphate (DIP) time series at the two EMS pelagic stations. Data shown is from the top 200 m although bottom depths are 1450 and 832 m for THEMO2 and N800 stations, respectively. The sampling in 2018-2019 are from THEMO2 station (Ben-Ezra et al., 2021) while from 2020 onwards the sampling are from station N800. The sampling depths are marked as dots, contouring was obtained using ODV weighted average gridding set for 60 permille x scale-length and 40 permille y scale-length.

Samples were collected from the RV Mediterranean Explorer using a rosette with 12 Niskin bottles. Physical data was collected using a Sea-Bird Electronics (SBE19V plus) CTD profiler equipped with conductivity, temperature, in-situ fluorescence chlorophyll (Chl), turbidity, photosynthetically active radiation (PAR), and for some cruises dissolved oxygen profiles. Samples were collected generally from about 10:00 until 13:00. Water samples were always taken from 12 depths from 2.5 to 200 m depth (2.5, 10, 25, 50, 75, 100, 125, 140, 150, 160, 180 and 200 m) in the water column. In some cases, the depths between 100 and 160 m were modified to account for the observed depth of the deep chlorophyll maximum (DCM; observed using the CTD Chl fluorescence sensor).

### 2.2. Nutrient sampling and analyses

Seawater samples for nutrient analyses NO_3_+NO_2_ (N+N), NH_4_, PO_4_ (DIP), and Silica (SiO_4_) were collected in pre-washed 500 ml bottles (1 time 10% HCL then washed 3 times with distilled water followed by once using Milli-Q Synergy double distilled water (MQ) and washed 3 times with sample water. The nutrient samples were immediately filtered using sterile Nalgene filters (0.22 µm). The Nalgene filters were first washed (twice) with the sample seawater and the remaining water was then filtered and transferred into triplicate 50 ml Falcon tubes. The fresh samples were stored at 4 °C, and replicate samples were frozen in a –20 °C freezer. The fresh samples were analyzed within ∼24 hours at the Sdot-Yam Marine Station with a SEAL Auto Analyzer 3 which was used to determine the nutrient concentrations. Frozen samples were used for potential backup for dissolved nutrient analysis if something went wrong with an analysis the day after the cruise. Sample preservation experiments have shown that filtered frozen samples were not significantly different from fresh filtered samples but were significantly different from unfiltered frozen samples (Ben-Ezra et al. 2023).

#### 2.2.1. Chemical analyses

NH_4_ was analyzed using an automated orthophthalaldehyde fluorescence method (SEAL Analytical 2011a). To avoid cross-contamination from analytical reagents, at the low concentrations of NH_4_ found in the EMS, NH_4_ samples were analyzed first and on their own. They were allowed to reach room temperature before the run. To minimize NH_4_ contamination from the air, the falcon tubes containing samples were only opened from the tray just before the automated sampler took the sample. The N+N analysis method uses Cd reduction to nitrite and diazo dye. SiO_4_ was run together with N+N by molybdate blue in the presence of oxalic acid (SEAL Analytical 2011b; c). To measure DIP a modification of the molybdate blue determination (Murphy and Riley 1962) was used. A 100 cm long flow cell (LWCC) was used as the detection device for the ultra-low concentrations of DIP found in the area. The baseline water used for the NH_4_, N+N, and SiO_4_ methods was fresh Milli-Q while the baseline for DIP was aged surface seawater from August 2020. This is due to issues of minor contamination (∼20 nM of DIP) of the MQ water (Ben Ezra et al. 2021).

### 2.3. Flow cytometry

To identify and quantify the predominant groups of pico and nanophytoplankton, triplicate seawater samples (2 ml each) were collected from various depths and placed in cryo-vials (SORFA Life Science). To preserve these samples, 5 μl ml^−1^ of 25% glutaraldehyde (Sigma) was added. The vials were then subjected to a 10-minute incubation in darkness at room temperature, flash frozen using liquid nitrogen, and stored at –80 °C. Prior to analysis, the samples were thawed under dark conditions at room temperature. For analyses, each sample underwent two runs on a BD Canto II flow cytometer, with the use of 2 μm diameter fluorescent beads (Polysciences, Warminster, PA, USA) as a size and fluorescence standard. In the initial run, three types of phytoplankton cells – *Prochlorococcus*, *Synechococcus*, and picoeukaryotes—were identified based on their intrinsic auto-fluorescence. These cells were distinguished by their cell chlorophyll (Ex482nm/Em676nm, PerCP channel), phycoerythrin fluorescence (Ex564nm/Em574nm), and cell size (forward scatter). Prior to the second run, the samples were stained with SYBR Green I (Molecular Probes/Thermo-Fisher) to facilitate counting, with detection at Ex494nm/Em520nm (FITC channel). This allowed for the quantification of the total microbial population (0.5-3 µm), encompassing the pico and nanophytoplankton, heterotrophic bacteria, and archaea. Additionally, it enabled the differentiation between cells with high or low DNA content. Data analyses were conducted using FlowJo software. Flow rates were consistently monitored during each running session by weighing tubes filled with double-distilled water, and the counts of the standard beads were employed to confirm a consistent flow rate. Biomass calculations were performed for each of the identified groups individually. The number of cells per ml were multiplied by 53, 175 or 2100 fgC cell^−1^ for *Prochlorococcus, Synechococcus*, and picoeukaryotes, respectively (Campbell 2001).

### 2.4. Wavelength dispersive XRF (WDXRF) analyses

Seawater samples for XRF analysis were filtered onto 47 mm polycarbonate filters with a nominal pore size of 0.45 μm. Triplicate samples of 1 L were collected and measured from each of three sample depths, 25, 100 and 180 m. After filtration, excess salt was removed by rinsing filters with a few ml of MQ water. After filtration the filters are placed in Petri dishes and allowed to air dry for up to 1 h. The dry filters were then stored at room temperature until they were sent to University of Bergen, Norway for analysis. Triplicate blank filters from each box were analysed, obtained by filtering 500 ml of prefiltered MQ water to remove background concentration from filters.

Elements were analysed using a fixed, optimized goniometer setting. Each element was measured until the analytical error due to relative statistical error was lower than 0.3%, but not for longer than 30 s. All WDXRF results are based on the intensities (kilocounts per second – kcps), and the analysis time for a sample was ∼12 min. The instrument measures total amount of the elements and does not distinguish between different chemical forms. Details of the calibration, machine settings and LoD are given in Paulino et al. (2013).

### 2.5. Satellite data

Time series of sea surface Chl concentrations and temperatures were extracted by averaging satellite data over the study area. The satellite dataset consisted of 1 km daily Level 3 products extracted from Copernicus Marine Environment Monitoring Service (CMEMS, https://marine.copernicus.eu/; D’Alimonte et al., 2003; Volpe et al., 2018). The CMEMS Chl product uses merged remote sensing reluctance data from multiple available ocean color sensors (MODIS-AQUA,NOAA20-VIIRS, NPP-VIIRS and Sentinel 3A-OLCI), produced by the Plymouth Marine Laboratory (PML) using the OC-CCI processor version. The Chl retrieval algorithm specifically tuned for the Mediterranean region by using the MedOC4 algorithm 20 (Volpe et al. 2019) for Case I waters, and the AD4 algorithm (Berthon and Zibordi 2004) for Case II waters.

## 3. Results

### 3.1. Overview of what data is being presented

In this section we describe in detail the data from January 2020 until June 2022. We then analyse and discuss this data in the context of previously published water column data from the THEMO2 site (Ben Ezra et al. 2021; Reich et al. 2022). For simplicity’s sake, both the previously published data and the new data are plotted together on the same 4.5 yearlong Ocean Data View (ODV; Schlitzer, Reiner, Ocean Data View, https://odv.awi.de, 2023) plots. Plotting data from THEMO2 and N800, although geographically quite close, transcends mere logistical practicality. Both stations lie offshore the Israeli coastal shelf, sharing exposure to similar regional processes and environmental drivers. Despite the depth difference, both stations are experiencing comparable light regimes and surface-driven influences and the lack of sediment-driven influences, when discussing photic zone processes. Furthermore, THEMO2’s established dataset, featured in previous publications, serves as a well-studied reference point for N800.

### 3.2. Physical characteristics of the water column during the sampling period

The temperature profiles in the upper 200 m showed seasonal changes with a well-mixed water column from January 2020 through March 2020 which was slightly cooler (by an average of 1.2 °C) than the winter of 2018 and 2019 (see Figure 9). The water column became seasonally stratified starting in April 2020. This surface stratified layer became deeper and warmer throughout the summer. Mixing had occurred by the first sampling in February 2021. By April 2021, the water column started again to be seasonally stratified. We did not sample between August 2021 and December 2021 (which was immediately after a major storm called Storm Carmel) at which time the water column showed only the permanent thermocline (presumed mixed) and remained so through the next 2 samplings in February and March 2022. Changes in salinity were consistent with temperature, with increased salinity (up to 39.8 psu) observed in the seasonally stratified upper water during the normal (for this region) hot dry summer and relatively low salinity (38.8 psu) modified Atlantic water immediately below the seasonally stratified layer. The well mixed water column in the winter of 2022 was unusual in having lower temperature and salinity (15.1-17.2 °C and 38.8-39.2 psu, respectively) compared with previous winters (17.6-19.2 °C and 38.6-39.2 psu).

**Table 1.** Analytical performance for nutrient analyses. The precision for DIP, N+N, NH4 and SiO_4_ was 2 x standard deviation of 6 replicate samples. The limit of detection (LoD) for inorganic nutrient analysis was 3 x the standard deviation of the blanks (5 replicates; Ben Ezra et al., 2021).

| Parameter measured | Range [nM] | Precision [nM] | LoD [nM] | Detector device used |
| --- | --- | --- | --- | --- |
| DIP | 0-125 | 2 | 6 | 100cm WPI long flowcell |
| N+N | 0-1000 | 34 | 13 | 1 cm cell spectrophotometer |
| $\text{NH}_4$ | 0-500 | 6 | 5 | JASCO fluorometer |
| $\text{SiO}_4$ | 0-2000 | 7 | 11 | 1 cm cell spectrophotometer |

**Table 3:**
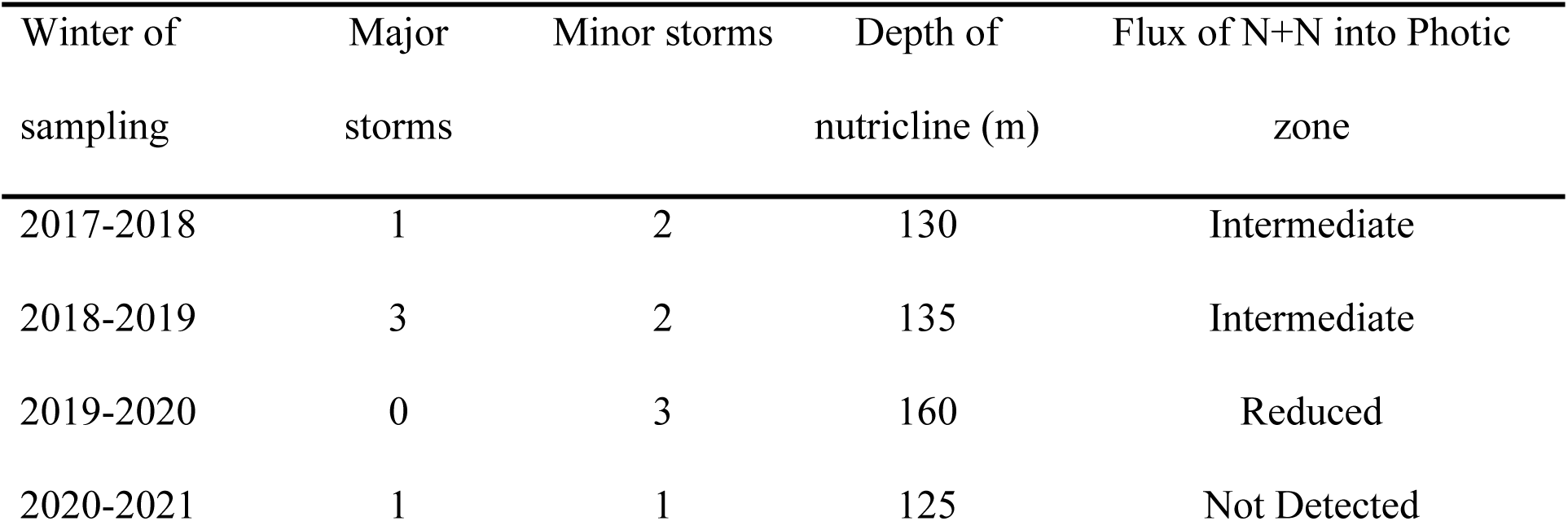
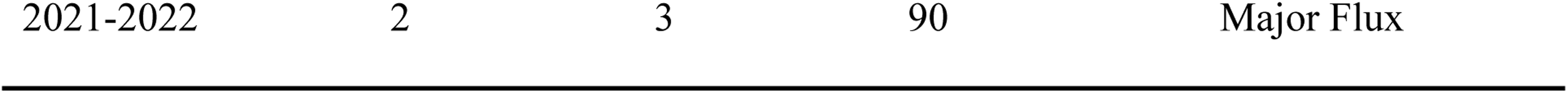
Major and minor storm count. The number and intensity of winter storms between 2017-2022 combined with the depth of the nutricline and flux of N+N into the photic zone.

### 3.3. Description of nutrient dynamics in the upper water column during the sampling period

The nutrient which showed the clearest seasonality was N+N with a peak in winter related to mixing of N+N from the top of the nitricline followed by a gradual decrease in N+N (down to below our LoD; see Table 2) during the spring/summer stratified season. From January to March 2020, the average N+N in the upper 100m was low (56 nM). The peak concentration was observed in April 2020 (average of 610 nM) which began to decrease by May 2020 (average of 480 nM). From June 2020 for the rest of the year, the upper waters below 2 m and above a nutricline of 125-150 m were depleted in N+N (<50 nM). Unusually, in the winter of 2020-2021 there was no increase in N+N in the upper water (average of 80 nM) and the upper water column remained depleted in N+N throughout 2021 (average of 75 nM). At the same time the nitracline gradually shallowed until, by the last sampling of the year in August 2021, the nutricline was observed at 100 m. From December 2021 through March 2022, the N+N in the water column was both homogeneous throughout the sampling depth and unusually high (Maximum of 1090 nM, average of 690 nM). By the last sampling in this study in June 2022 after seasonal stratification was established, the N+N had decreased somewhat particularly in the upper seasonally stratified waters (180 nM) but was still considerably higher than at similar periods in 2020 or 2021. Throughout the sampling period, DIP measurements remained at or below 10 nM and often below 5 nM except for occasional single data points in the photic zone which were higher than 10 nM but less than 25 nM. Occasionally higher values (29 and 48 nM) were measured in the surface waters (e.g. autumn 2020).

NH_4_ was also generally low throughout most of 2020 and 2021 (average of 32 nM). We measured higher values of NH_4_ in August 2021 (average of 245 nM) and in December 2021 (average of 187 nM) but the main increase occurred in February 2022 when NH_4_ values reached an average of 750 nM. By June 2022, the NH_4_ concentrations had decreased and were close to their limit of detection (5 nM; Table 2).

Dissolved silica (SiO_4_) in the photic zone typically ranged from 0.6-1.5 µM. The major change in silica concentrations was observed in the winter/spring of 2021-2022 when silica values dropped to ∼0.1 µM. By June 2022, silica had returned to ‘typical’ values in summer (∼1 µM) (see Figure 10 for a composite figure discussing the probable reasons for these changes in silica concentration).

Major element analyses of particulate matter were carried out during the period from April 2020 until January 2021. Particulate Fe:Si (Figure 5a) showed a very close linear relationship with a slope of 0.24 (µmole L^−1^ vs µmole L^−1^; R^2^ = 0.97) calculated from the July 2020 sample except for December 2020 when the particles were unusually Fe rich. This sampling had the highest particulate matter content estimated to be 30 µg SPM L^−1^ calculated by assuming that the measured Si in the water column had the typical Si content per unit mass measured in Saharan dust (Eijsink et al. 2000; Krom et al. 2016; Stockdale et al. 2016). The plot of particulate O:Si (Figure 5b) shows a similar tight linear relationship with a slope of 1.85 (µmoleO L^−1^ vs µmoleSi L^−1^; R^2^ = 0.97) calculated from the July 2020 sample. There was a similar very minor increase in O:Si ratio in December 2020 and January 2021 compared to other months (Figure 5b; Stockdale et al., 2016).

**Figure 4:**
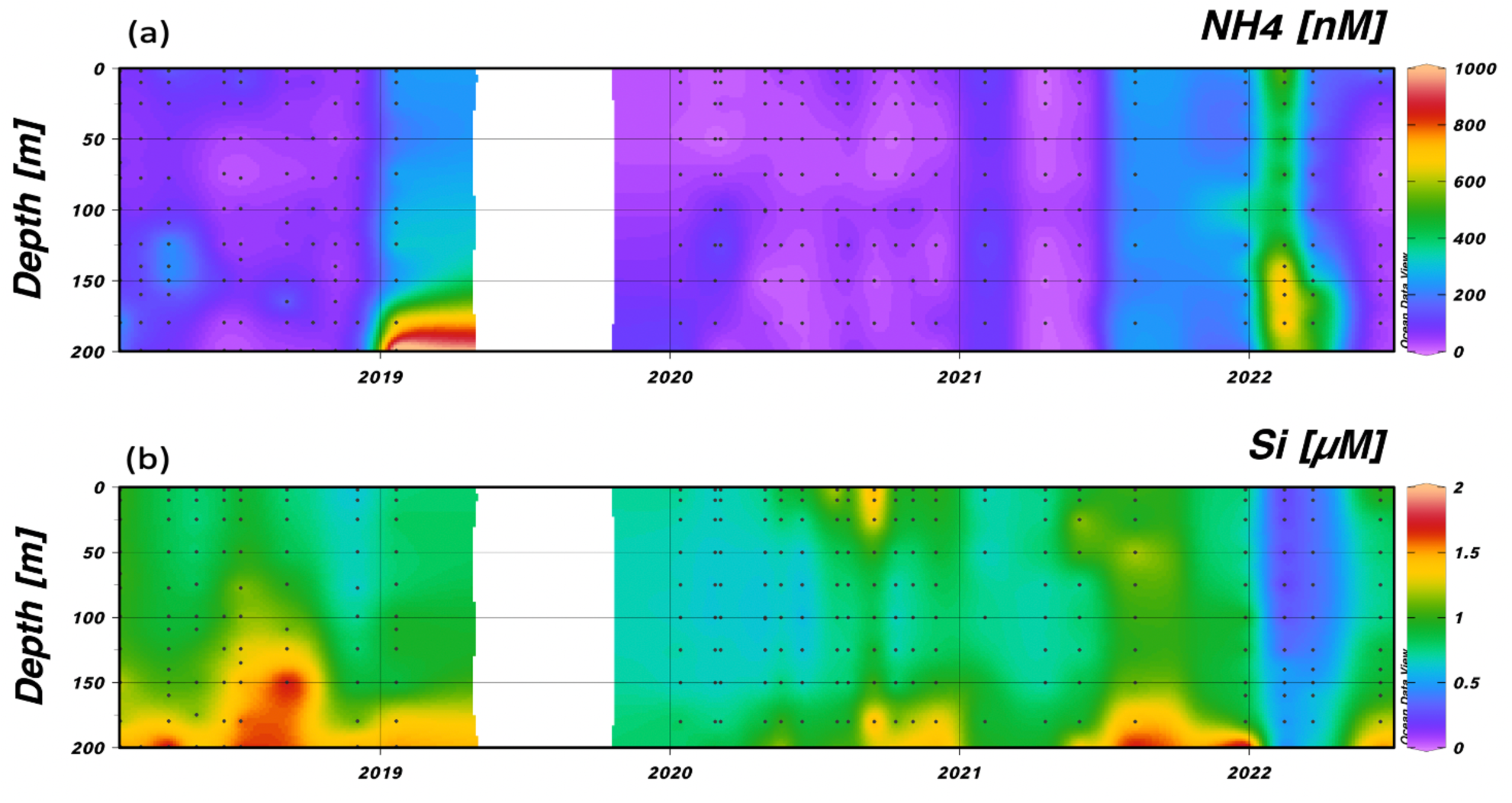
(a) Ammonium (NH_4_; nM) and (b) dissolved Silica (Si; µM) time-series at the two EMS pelagic stations. Data shown is from the top 200 m although bottom depths are 1450 and 832 m for THEMO2 and N800 stations, respectively. The sampling in 2018-2019 is from THEMO2 station (Ben-Ezra et al., 2021) while from 2020 onwards the data are from station N800. The sampling depths are marked as dots, contouring was obtained using ODV weighted average gridding set for 35 permille x scale-length and 140 permille y scale-length.

**Figure 5:**
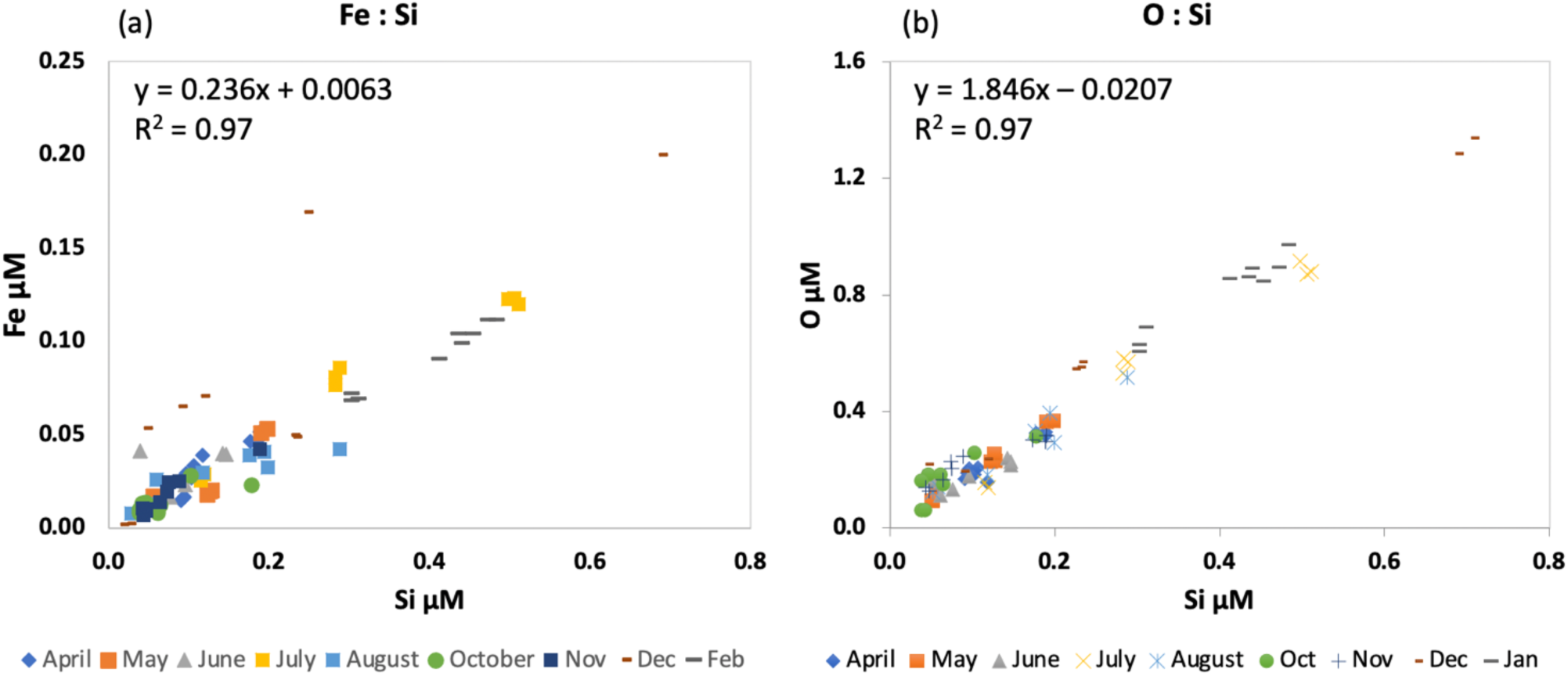
(a) Particulate Fe vs particulate Si and (b) particulate O vs particulate Si sampled at 25, 100 and 180 m in the upper water column from April 2020 to January 2021. The samples were determined using thin film WDXRF.

### 3.4. Description of phytoplankton biomass from winter 2021 to winter 2023 in the upper layer (0-200 m)

The pico and nanophytoplankton (*Synechococcus*, *Prochlorococcus* and picoeukaryotes) as determined by flow cytometry and calculated as biomass distinctly differed in their vertical distribution within the upper 200 m. The annual study (monthly sampling) from 2018 at THEMO2, (Reich et al., 2022; Figure 6) showed that *Synechococcus* was mainly present during winter mixing and tended to predominate in the upper 50 m of the water column. *Synechococcus* was also found during summer stratification mainly in the upper water layers but with reduced biomass. By contrast *Prochlorococcus* was found predominantly towards the base of the photic zone just above the nutricline and in the greatest biomass when the water column was seasonally stratified. The highest biomass of phytoplankton were picoeukaryotes which were found mainly from 50-200 m during winter mixing and at the DCM for the rest of the year.

**Figure 6:**
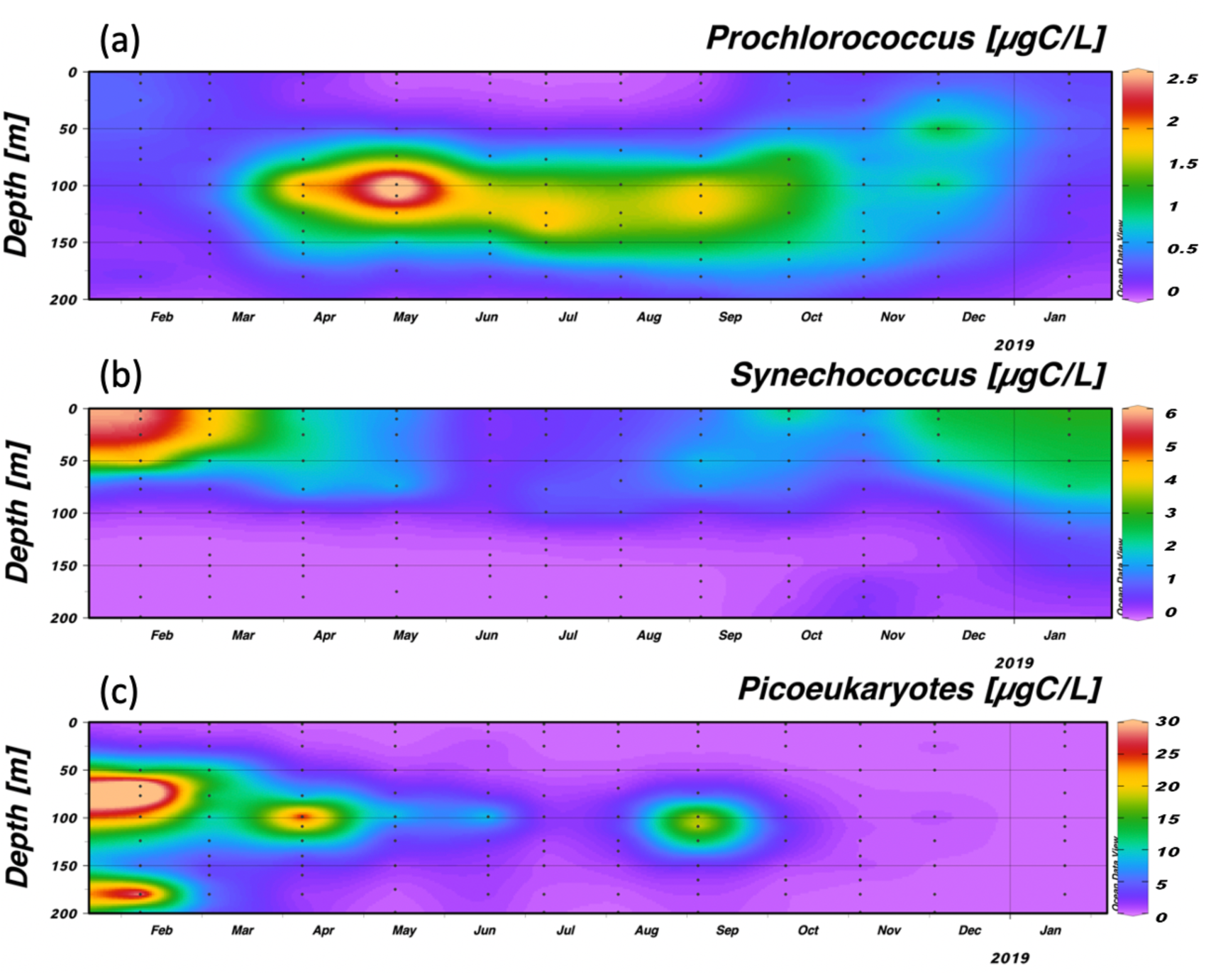
Phytoplankton biomass calculated from flow cytometry data for (a) Prochlorococcus; (b) Synechococcus and (c) picoeukaryotes. Data from February 2018 until January 2019 from the THEMO2 station was calculated from data in Reich et al. (2022). Data shown is from the top 200 m. The sampling depths are marked as dots, contouring was obtained using ODV weighted average gridding set for 40 permille x scale-length and 60 permille y scale-length.

Here we measured the distribution of these three groups during 2021 and 2022. During the period of nutrient depletion of 2021, the depth distribution was similar to that of 2018 but the phytoplankton had much lower biomass (Figure 7). Thus, *Synechococcus* was generally more abundant in the upper 50 m, while picoeukaryotes and *Prochlorococcus* were found in deeper waters near the DCM/nutricline. This pattern changed dramatically after Storm Carmel. By far the dominant species measured by FCM were now picoeukaryotes and these were found throughout the water column. *Synechococcus* was found in December 2021 but not in February 2022 while the converse was true for *Prochlorococcus* which was found March 2022 but not in December 2021. By June 2022, the depth distribution of pico and nannoplankton returned to the pattern observed in 2018 though the biomass was considerably higher.

**Figure 7:**
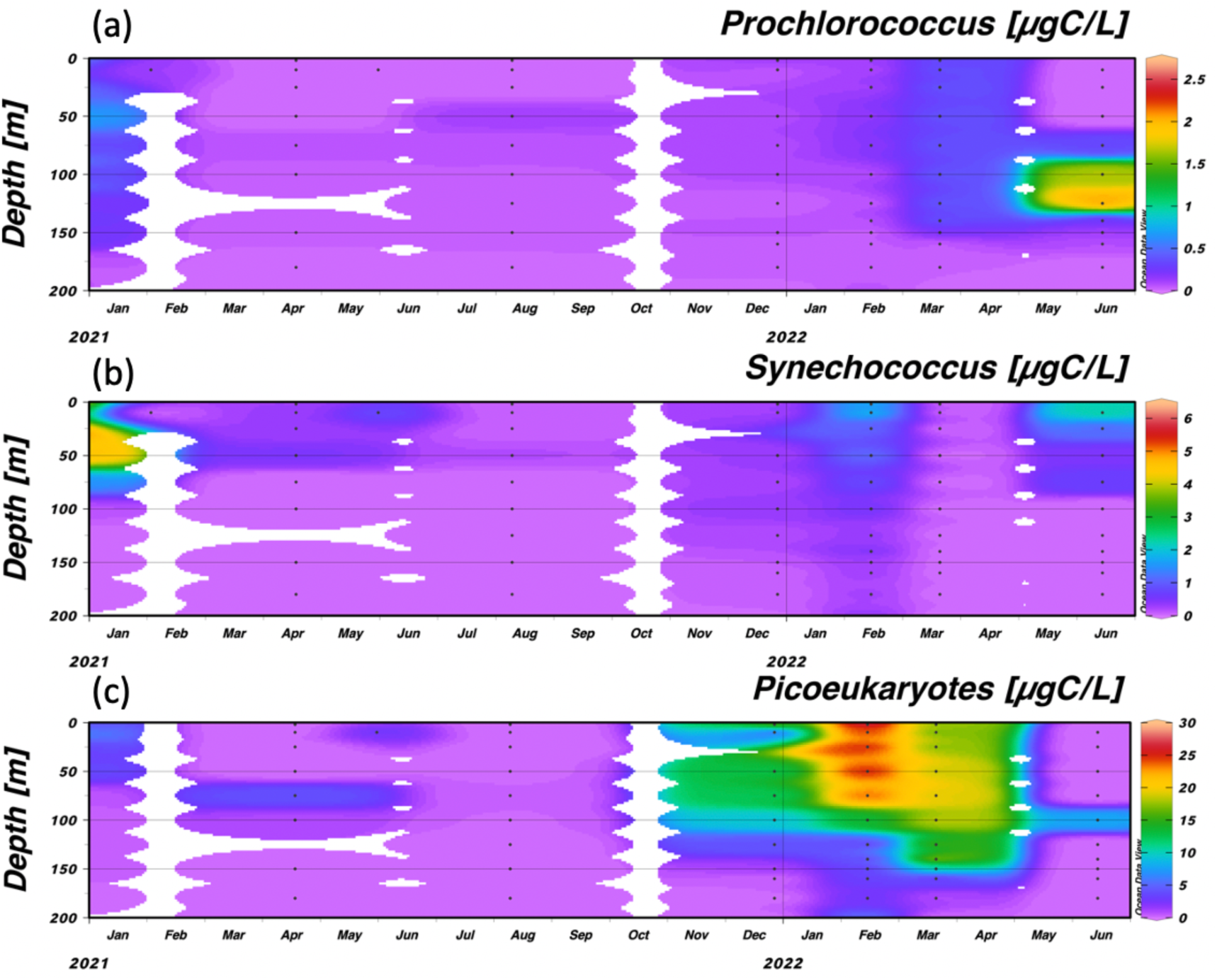
Phytoplankton biomass calculated from flow cytometry data for (a) Prochlorococcus; (b) Synechococcus and (c) picoeukaryotes from March 2021 until June 2022 from station N800. Data shown is from the top 200 m. The sampling depths are marked as dots, contouring was obtained using ODV weighted average gridding set for 40 permille x scale-length and 60 permille y scale-length.

The total biomass of the three characterized pico and nanophytoplankton, as determined from flow cytometry measurements during 2018, averaged 280 ± 90 μgC m^−2^ with a general seasonal trend (Figure 8). Higher biomass was typically measured during the mixed water column period – winter (average of 247 ± 55 μgC m^−2^ in February 2018 & December/January 2018-2019) than during the thermally stratified summer and fall (average of 91 ± 63 μgC m^−2^; Reich et al., 2022). There was a similar biomass and seasonal distribution pattern in 2020 (Steindler L. unpublished data). By April 2021 the period of maximum nutrient depletion, the total biomass had decreased to 80 μgC m^−2^ and further decreased to 20 μgC m^−2^ by August 2021. Phytoplankton biomass immediately increased 3 days after Storm Carmel to 1050 μgC m^−2^ and growth continued with biomass reaching a maximum of 2400 μgC m^−2^ in February 2022. By March 2022, biomass had declined to 1747 μgC m^−2^ and reached a relatively low of 280 μgC m^−2^ by June 2022 when thermal stratification was pronounced. The calculated autotrophic biomass was less than 20% of the maximum value recorded in February 2022, and similar to the biomass recorded during mid-summer 2018 and 2020. At 10 m depth there was a systematic decrease in phytoplankton biomass from December 2020 to August 2021 in the depleted period followed by a drastic increase after Storm Carmel in December 2021 (see Fig S2).

**Figure 8:**
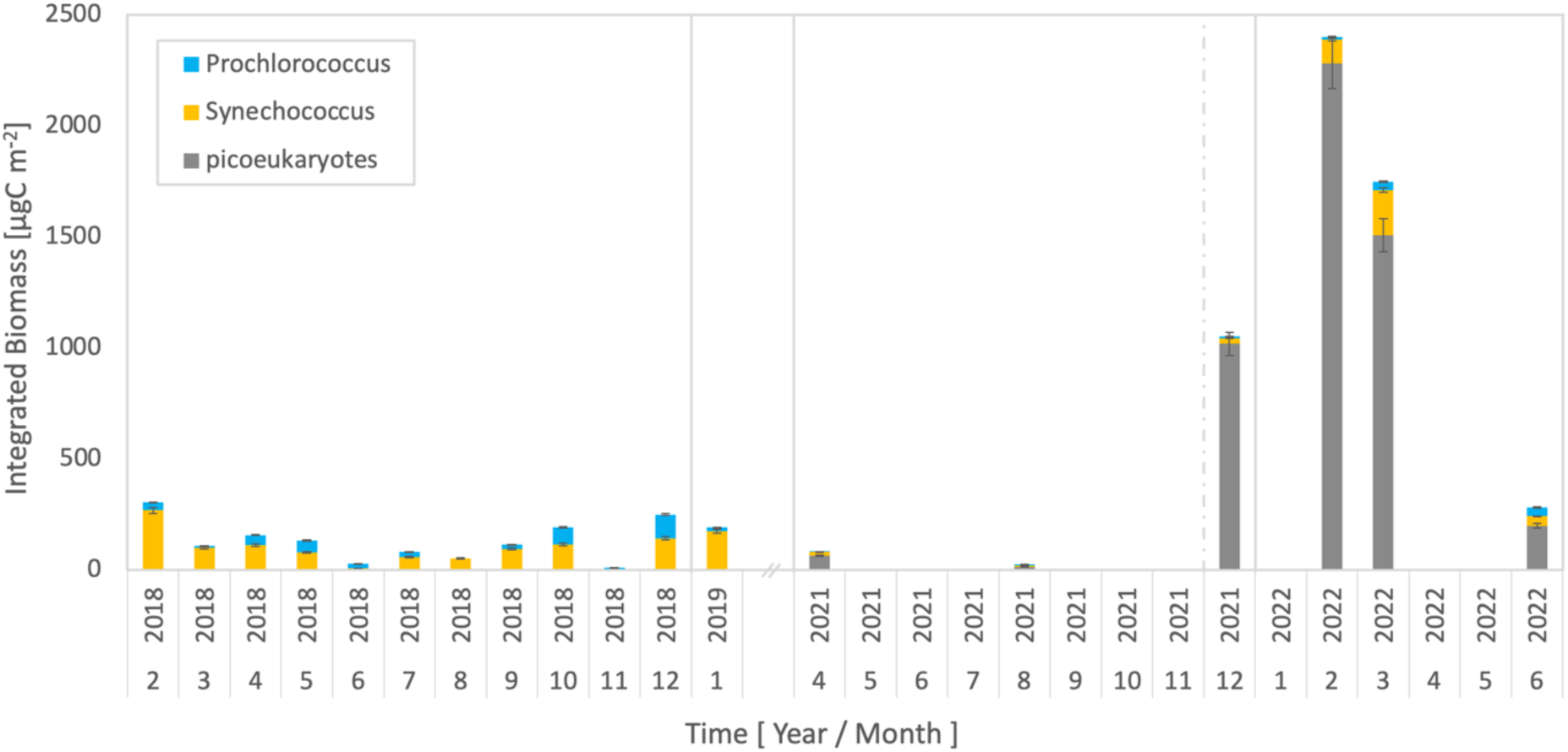
Temporal changes in integrated biomass (surface to 100 m depth) of the measured pico and nanophytoplankton based on cell count measurements in 2018 (THEMO2) and at the time series site (N800) from April 2021 onwards. Note that the time axis is cut off during 2019-2021.

### 3.5. Remote sensing measurements of the upper water column at the 800m station

Figure 9 shows the annual surface temperature changes, determined from satellite measurements, at the N800 time-series station. Using this data alone, while there is a clear annual signal in temperature, there is no observable systematic increase in the temperature signal from 2018 until 2022 although the winter of 2021 was warmer (average of 20.3 °C) than other winters while 2022 was slightly colder (average of 19.5 °C). On the X-axis is marked the date of major and minor winter storms including the cluster of major storms in the winter of 2021-2022 which start with Storm Carmel on December 17-24, 2021 and continued through the winter. The fluctuations in surface Chl sometimes are adjacent to such storms (2019 and 2022) and at other times the Chl peaks are not directly related to storms.

### 3.6. The number and magnitude of winter storms during the sampling period

An important driver of the nutrient and phytoplankton dynamics are winter storms which in the EMS mix nutrients into the photic zone followed commonly by periods of calm and sunlight that allow rapid biogeochemical responses. Accordingly, we posit that two key parameters in determining the magnitude of the flux of nutrients into the photic zone in winter are the number and intensity of winter storms and the depth of the nutricline in the period before the first storm in winter (Table 3). Major winter storms were defined by a Beaufort scale wind of 9 or 10 lasting for a period of >24h and minor winter storms were defined by a Beaufort scale wind of 8 for >24h (“Windguru Station 2049,” https://www.windguru.cz/885332).

During 2017-2018 1 major and 2 minor storms occurred while during 2018-2019 3 major and 2 minor storms were recorded (Table 3). For both years, the nutricline depth was 130 m prior to the start of winter. In the winter of 2019-2020, there were no major storms, and only 3 minor storms and the nutricline was 160 m. In 2020-2021, despite the observation that there was 1 major storm and 2 minor storms, and the nutricline was at 125 m at the beginning of the year, there was no evidence of any ‘new’ N+N mixed into the photic zone. It is possible with the timing of the sampling that we might have missed a small influx of N+N but there certainly was much less than in 2017-18 or 2018-19 or even 2019-20.

The winter with the major flux of nutrients into the photic zone was 2021-2022 in which there were 2 major storms (one of which was called ‘Storm Carmel’) and 4 later minor storms with the initial nutricline at 90 m. This winter was also colder than previous winters (Figure 9).

**Figure 9:**
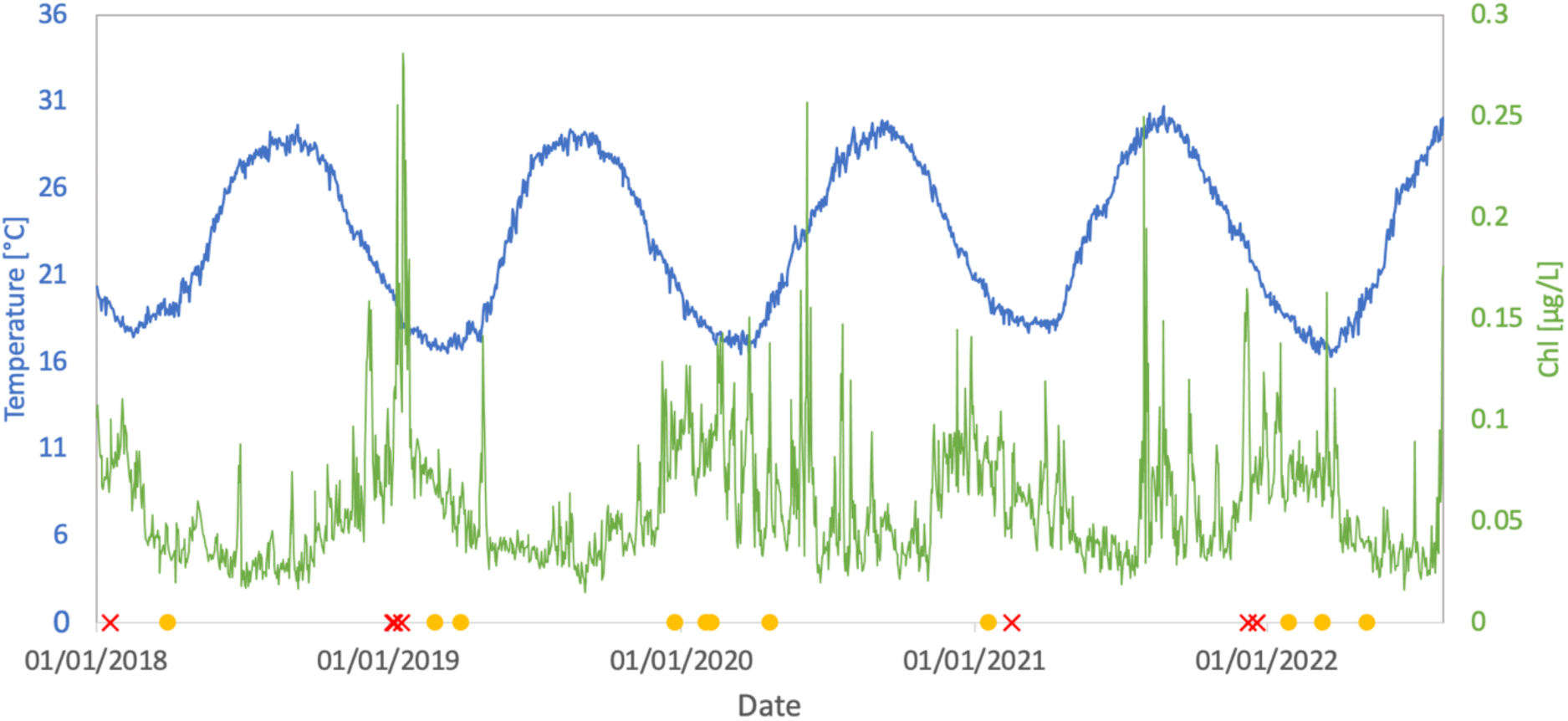
Seasonal changes in satellite-derived temperature (blue) and Chlorophyll-a (green) at the N800 station over a period of 4.5 Years (2018-2022). The timing of major (red x) and minor storms (yellow circles) are marked on the X-axis (see Table 3).

**Figure 10:**
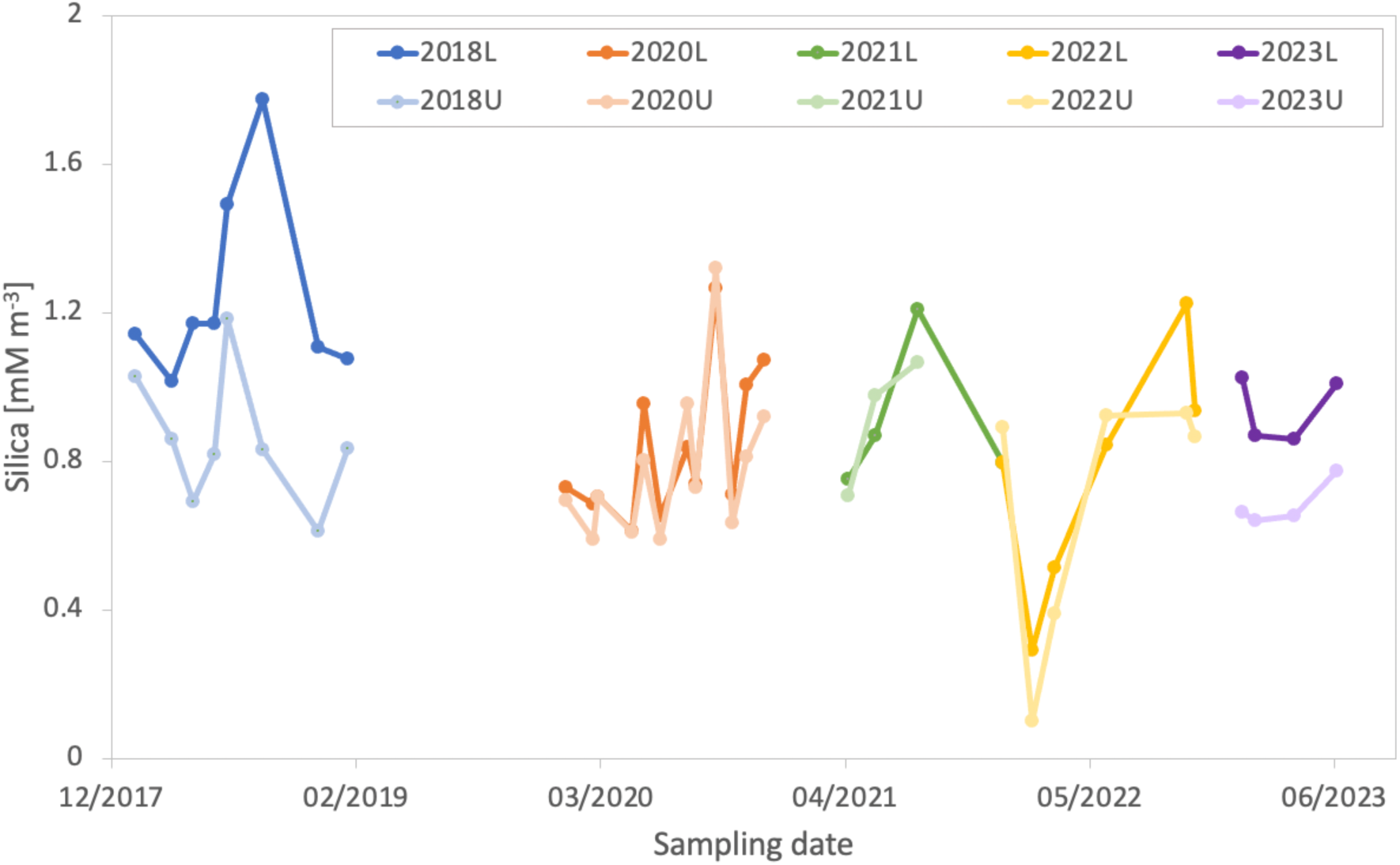
Plot of average dissolved Silica in the water column calculated over the upper 100m (years L) and from 100 m to the top of the nutricline (years U) for each sampling year from 2018 to 2023.

## 4. Discussion

### 4.1. Nutrient dynamics and phytoplankton biomass define normal, depleted and mesotrophic status of the ecosystem

Nutrient dynamics and resulting autotrophic picophytoplankton biomass changes show clear annual and interannual variability in the ultra-oligotrophic EMS (Kress and Herut, 2001, Ben Ezra et al., 2021; Reich et al., 2022). To further explore the drivers influencing nutrient and phytoplankton dynamics we have established a time-series station off the Israeli coast (at station N800). Our initial measurements are in the spirit of the BATS station off Bermuda in the N. Atlantic gyre (Cavender-Bares et al. 2001; Steinberg et al. 2001; Lomas et al. 2013; Djaoudi et al. 2018a) and have so far been running for 5 years.

Based on our results, three ecosystem states are recognised (Normal, Depleted and Mesotrophic). These ecosystem states correspond to potential future conditions in the EMS under climate change. The measured changes in N+N provide the clearest marker for these fundamental changes in nutrient availability and subsequent phytoplankton response. As is typical of such ocean systems, N+N accumulates below the photic zone during the year. As autochthonous organic matter drops out of the upper layers, a fraction is then respired, releasing reduced N (NH_4_) and DON. In addition, zooplankton and other microorganisms often migrate out of the photic zone diurnally and excrete reduced N below the photic zone. This NH_4_ is then nitrified as has been shown previously particularly at the top of the nutricline in the EMS (Krom et al. 2005b; Pujo-Pay et al. 2011; Ben Ezra et al. 2021) and nitrate accumulates. When N+N is mixed into the photic zone during winter mixing, initially only part of the total input is taken up rapidly by phytoplankton (Krom et al. 1992) while the remainder accumulates in the water column. This partial uptake of N+N is shown both by the residual N+N in the water column in winter (Figure 3) and its isotopic composition which is characteristically heavy (Emeis et al. 2010). At the same time, isotopically light PON and TDN are formed through growth of phytoplankton and its subsequent breakdown. The magnitude of N+N measured in the photic zone is a balance of the flux of N+N into the photic zone and the remainder after phytoplankton uptake (Krom et al. 1992; Ben Ezra et al. 2021).

During the period which we identify as ‘Normal’ conditions (2018-2019) the average concentration of N+N in the upper 100 m of the water column, was intermediate with 470 nM (February/March 2018) and 230 nM (December 2018/January 2019; Figure 3; Ben-Ezra et al., 2021) in winter. These values decreased to ∼20 nM in September & October 2018. These are similar to the range of N+N values observed in Levantine surface water by Kress and Herut (2001). As is discussed below, DIP remained at <10 nM (average of 4 nM) throughout this period and Silica was low in winter and gradually increased through the period of seasonal stratification. The vertical distribution of pico and nanophytoplankton was characteristic to the southeastern Levantine basin. During winter mixing, both *Synechococcus* and picoeukaryotes were evenly distributed throughout the water column. As stratification set in, *Synechococcus* primarily resided in the upper waters, coinciding with nitrate accumulation in the photic zone. In contrast, *Prochlorococcus* remained scarce in winter but increased near the nutricline during stratification (Reich et al., 2022). This pattern aligns with established knowledge of their ecological preferences. Furthermore, picoeukaryotes exhibited a similar shift, transitioning from a uniform winter distribution to concentrating around the DCM just above the nutricline during stratification. These findings correspond with the ultra-oligotrophic nature of the southeastern Levantine basin, exhibiting a total biomass of up to 670 µgC m⁻² in the top 100 m during 2017-2018, with a slight decrease in summer. They fall within the lower spectrum of what Denis et al. (2010) observed across the EMS, where they reported peak values of 2200 µgC m⁻² for total pico and nano eukaryotes and ∼600 µgC m⁻² for *Synechococcus* alone.

The minimal flux of nutrients mixed into the photic zone during the nutrient ‘Depleted’ period starting in the winter of 2020 and especially during 2021, coincide with the period with less winter storms than the rest of the study period (Table 3). N+N concentrations in the water column, below the top surface layer, decreased to <50 nM (i.e. below our LoD). Phosphate remained at or below detection limits while Silica was the same concentration in the upper 100 m as it was between 100 and 180 m (see below). During the nutrient depleted seasonal stratification in 2021, the depth distribution of pico and nanophytoplankton remained similar to that observed during the ‘normal’ period (Reich et al., 2022) with *Synechococcus* predominating in the upper water column and *Prochlorococcus* and picoeukaryotes found mainly at the DCM which was just above the nutricline. However, the biomass of phytoplankton during this period of nutrient depletion was much lower, reaching a minimum of 23 µgC m^−2^ in August 2021 while the concentration measured at 10 m was <3 μgC L^−1^ down from values of >10 μgC L^−1^ in September 2020. These values were much lower compared to the ‘normal’ period of 2018-2019 (Reich et al., 2022, Denis et al., 2010).

Storm Carmel (December 2021), which was defined as a major storm (Table 3), stimulated a major increase in the N+N concentration and based on the observed and inferred phytoplankton biomass, we describe this period of temporarily “Mesotrophic”. The N+N increase was higher than any result previously measured in the area (Max 1090 nM; Kress and Herut, 2001). This increase remained during the February and March 2022 samplings in part due to the continued stormy and cold weather that winter (Table 3, Figure 9). Subsequently, there was a major increase in NH_4_ (and probably DON) in the water column, likely due to recycling from the increased biomass. The increase in NH_4_ in the water column corresponded to the only period when *Prochlorococcus* was found throughout the water column (Figure 7). It is generally observed that *Prochlorococcus* uses preferentially NH_4_ for its N source and shows a limited ability to use nitrate (Masuda et al. 2023). Surprisingly, DIP did not increase even in the December 2021 sampling immediately after Storm Carmel. The reason for this observation is discussed below. After Storm Carmel, the considerable increase in pico and nanophytoplankton biomass (reaching a maximum 2400 µgC m^−2^) together with the population being entirely picoeukaryote-dominated while *Synechococcus* and *Prochlorococcus* abundance decreased considerably (down to 3% of the population), posing strong evidence for the ecosystem shift that may be caused by strong winter storms. Another demonstration of this shift is the dramatic decrease in silica (∼90% to ∼0.1 µM in February and March 2022), for the only time in the sampling period. The declining concentrations of Silica is likely due to uptake by silicious eukaryotic phytoplankton such as diatoms and silicoflagellates (Krause et al. 2009). In general, the only period in which there are a moderate number of large eukaryotes including diatoms in the water column of the EMS is in winter when nutrients are at a maximum (Ignatiades et al. 2002; Keuter et al. 2022). During ‘Normal’ winters the silica concentration decreases by ∼10% (see Figure 10). Although we did not measure the microplankton eukaryote biomass, it is likely based in part of the observed silica decrease, that during the winter of 2021-2022, larger eukaryotes including diatoms were present in considerably higher numbers than is normal and maybe dominated the autotrophic community temporarily.

The period after the winter 2022 represented a period of nutrient relaxation and the system decreased back to its ‘normal’ status. The concentration of N+N in the water column decreased by June 2022 to ∼50 nM. Pico and nanophytoplankton biomass was reduced and their depth distribution returned to that observed in 2018. All of these parameters decreased back to the lower values measured in 2018 (Ben-Ezra et al., 2021, Reich et al., 2022).

The observed pattern of NH_4_ was less systematic than N+N with NH_4_ being observed often as single elevated values. This is likely because NH_4_ is produced by grazing and/or phytoplankton respiration by bacteria but it is also rapidly consumed as the most bioavailable N compound (Berthelot et al. 2021). As a result, there are short term transient peaks of NH_4_ throughout the photic zone. Additionally, atmospheric inputs occasionally introduce N sources including NH_4_ resulting in higher concentrations of N in the surface waters where the typical combination of low nutrients and high irradiance inhibit high phytoplankton concentrations (Carbo et al. 2005; Theodosi et al. 2019).

### 4.2. Processes controlling phosphate distribution in the water column

DIP was measured for the first time using improved sampling techniques and high sensitivity analysis over a period of 4.5 years in the photic zone of the EMS, (Ben Ezra et al., 2021; Figure 3). The extremely low concentrations of DIP (almost always <5 nM) over different seasons is comparable with low POP measured in 2020 (average of 15 nM; maximum 70 nM) at N800, and DOP measured previously in the region (∼50 nM; Ben Ezra et al., 2021; Krom et al., 2005b) and average 40 nM (Pujo-Pay et al. 2011). The Fe vs. P plot (Figure S2) was typically at or lower than 1.5Fe:1P. This confirms that by far the largest fraction of particulate P in the water column was biological i.e. not associated with detrital Fe where the ratio of Fe:P in Saharan dust is typically 26:1 (Stockdale et al., 2016).

These results for DIP are similar to previous studies carried out in mid-summer during periods of seasonal stratification in the EMS (Krom et al. 2005b; Pujo-Pay et al. 2011; Djaoudi et al. 2018b), when all nutrient concentrations tend to be low (Kress and Herut 2001). However, the concentration of DIP in surface water was also low in winter when the concentration of N+N and other nutrients were relatively high (Zohary and Robarts 1998; Kress and Herut 2001; Ben Ezra et al. 2021).

In this study, even within 3 days of the end of the major Storm Carmel, when the concentration of N+N in the water column increased dramatically to 1090 nM, the average DIP in the upper 125 m averaged 4.1 nM. It is known that in this chronically nutrient depleted system, cyanobacteria, mainly *Synechococcus* and *Prochlorococcus,* are an important component of the phytoplankton community (Casotti et al., 2003; Denis et al., 2010; Reich et al., 2022; Figure 8 and 9) while SAR11 is the most abundant heterotrophs in this ultra-oligotrophic system (Haber et al. 2022). Previous studies in the P depleted N. Atlantic gyre have shown that *Prochlorococcus* and SAR11 are adapted to take up DIP very efficiently whenever these microorganisms come into contact with an increased pulse or patch of DIP (Zubkov et al. 2007, 2015). Kamennaya et al., 2020 showed that *Synechococcus* as well as *Prochlorococcus* and SAR11 will remove DIP to concentration levels of 10^−15^ M and store it as bioavailable inorganic P (PPi) within their periplasm. We hypothesize that this mechanism operates in these groups (and possibly others) in the EMS and that this is the reason for the consistently low levels of DIP found in the photic zone.

Using the data in this study it is possible to calculate the amount of missing DIP in the water column after deep winter mixing. The December 2021 sampling was 3 days after the end of Storm Carmel. The measured N+N in the water above the nutricline in December 2021 was 96 mmolesN m^−2^ compared with 1.6 mmolesN m^−2^ in August 2021 (Table S2). If we assume that the increase in N+N came from the upper layers of the nitricline, then it is calculated that it came from the top 100 m of the nutricline as measured in August 2021 i.e. from 100 m to 200 m. That depth interval contained 94.4 mmolesN m^−2^. That same depth interval (100-200 m) in August 2021 contained 2185 µmolesP m^−2^ of DIP. If we further assume that during Storm Carmel an additional 2185 µmolesP m^−2^ of DIP was added to the existing 459 µmolesP m^−2^ present in the photic zone during August 2021 then the increase in concentration present in the photic zone in December 2021 should have been 2644 µmolesP m^−2^. Yet, the amount measured was actually 817 µmolesP m^−2^ i.e. there was 1827 µmolesP m^−2^ missing. This converts to a calculated average of 13.6 nM of DIP over 100 m missing compared to the average measured concentration of 8.1 nM. If this missing DIP has been taken up by cyanobacteria, this calculation shows that it is 1.7 times the concentration measured in the water column.

Further evidence that there is a previously unmeasured reservoir of bioavailable DIP is provided by the observation that the concentration of DIP in EMS waters measured on unfiltered frozen samples (Krom et al. 1992; Zohary and Robarts 1998; Kress and Herut 2001) was typically twice the values obtained in this study. This would be expected if freezing and thawing cyanobacteria cells would release the DIP in the periplasm. In a study testing sample preservation procedures, in the EMS, Ben-Ezra et al. (2023) found that filtered fresh samples were in the range of 2-10 nM compared with unfiltered frozen samples 10-30 nM.

During the CYCLOPS Lagrangian addition experiment in May 2003, the dominant phytoplankton were cyanobacteria. It was observed that, following the addition of 18 tons of phosphate-enriched water, >80% of the DIP was taken up into the particulate P (microbial community) within a few hours of the Lagrangian addition, with no excess DIP remaining in the water column after 48 hours (Thingstad et al. 2005; Krom et al. 2005a). One consequence of this explanation is that soon after DIP is supplied to the cyanobacterial community, while it is no longer measured in the water column, it is still bioactive. A nutrient limitation experiment was carried out in CYCLOPS starting 3 days after the Lagrangian addition found that organisms which had been exposed to P acted as though they contained bioavailable P even though there was essentially no P in the water column (Zohary et al. 2005).

Although the measured concentration of DIP in the photic zone in the EMS was always low (<10 nM) it was not as low as the predicted threshold of DIP uptake of 10^−15^ M (Zubkov et al. 2007; Kamennaya et al. 2020). As is also found for NH_4_, the actual amount measured in the water column is a balance between the amount excreted by respiration and excretion processes in the water column and rapid uptake by cyanobacteria. This property of DIP in the water column has been observed previously in the P depleted N. Atlantic gyre by Zubkov et al. (2007), who showed that bioavailable P, as determined by ^32^P uptake, was somewhat different from measured DIP in the photic zone. It is thus suggested that many aspects of the P cycling in the EMS are similar to those observed in the P depleted N. Atlantic gyre (Reynolds et al., 2014, Wu et al., 2000)

### 4.3. Processes controlling silica distribution in the water column

In the EMS silicate increases gradually in the deep water from 6 µM in the S. Adriatic to a maximum of 12 µM in the southeastern Levantine basin (Krom et al. 2014b). However, that study showed that the entire increase could be accounted for by silica fluxing out of the underlying sediment from the breakdown of bioavailable and/or lithogenous particulate silica. The upper waters are effectively isolated from this deep reservoir which explains the low concentrations found in these waters of <1 µM (Crombet et al. 2011) and 0.6-1.8 µM (this study). During the stratified period of the year, Crombet et al., (2011) found no evidence of diatoms in June 2008 across the Levantine basin except for a small increase near the top of the nutricline in the core of the Cyprus eddy while Psarra et al. (2005) showed that in May 2002, there were only ∼0.1 diatom cells ml^−1^ and Fucoxanthin was ∼4% of the total pigment. The only time of year when diatoms become ∼10% of the total phytoplankton biomass was during winter mixing when total nutrients were at a maximum (Vidussi et al. 2001). During this study Reich et al. (2022) found higher fucoxanthin, which was interpreted as diatoms, only in January 2019.

In this study the measured Fe:Si ratio in 2020-2021 was within the typical range of Fe:Si measured from Saharan dust collected in the area (Eijsink et al. 2000; Krom et al. 2016). There was no evidence of any deviation from this single line as would be expected if there was any measurable biogenic silica except in January 2021 where the slope of Fe:Si showed a possible increase of 0.6% particulate Silica (SiO_2_) from the other samplings. There was a similar very minor increase in O:Si ratio in December 2020 and January 2021 compared to other months (Figure 5b).

In Figure 10 the average concentration of silica in the upper 100 m is compared with that from 100m to the top of the nutricline. It shows both an annual and interannual pattern which corresponds closely to the subdivision of our sampling periods into ‘Normal’, ‘Depleted’ and ‘Mesotrophic’. In all years the lowest value of silica was during winter mixing that was likely due to a combination of mixing of silica from the top of the nutricline and some uptake by diatom growth (Vidussi et al. 2001). In the ‘normal’ years of 2018, and the summer 2022 and 2023, the average silica concentration in the upper 100 m waters was slightly lower compared with the waters between 100 m and the nutricline. This suggests some uptake of silica in the upper water column, possibly by a minor population of diatoms, since silica is used in surface waters almost exclusively by diatoms (Krause et al. 2009). Intriguingly, for the depleted years 2020 and 2021, silica concentrations were uniform throughout the water column implying a total lack of diatoms. In the 4.5 years of this study, silica increased gradually from low values in winter to higher concentrations of silica throughout the summer. The almost constant Fe:Si ratio measured (0.24) is close to the value expected for dust, which is known to be incident in relatively large amounts in this area (Guerzoni et al. 1999; Ganor et al. 2010). The systematic increase in silica with time each year, but most clearly in 2020 and 2021, is likely to be due to dissolution of silica from weathering of inorganic particulate matter in the warm surface waters of the EMS. As noted above the only major decrease in silica was in winter 2021-2022 soon after Storm Carmel and the subsequent major increase in nutrients, which is interpreted as a diatom bloom.

### 4.4. Changes in nutrient flux and phytoplankton biomass as predictors of the effects of climate change on the EMS and similar systems

While the EMS has been identified as a location particularly vulnerable to climate change (Ozer et al. 2017), a period of 4.5 years is not sufficient to identify major changes in nutrient dynamics and phytoplankton productivity due to climate change. Nonetheless we suggest that the patterns observed between 2018 and 2022, are characteristic of the type of changes in nutrient supply to the photic zone predicted by the long-term changes in climate (Behrenfeld et al. 2006). Thus, compared with the ‘Normal’ nutrient patterns and phytoplankton distributions in 2018 and 2019 (Ben Ezra et al. 2021; Reich et al. 2022), 2020 and more severely 2021 represent a period of extreme nutrient depletion. The low fluxes of nutrients were the driver of the extremely low pico and nanophytoplankton biomass measured in the upper 100 m (2.9 µgC m^−2^; Figure 8). The summer and autumn period is always a period of reduced resource availability for the base of the food web in the EMS but this period of especially low nutrient availability we suggest would result in a more extreme period of climate driven ‘famine’ for the entire pelagic ecosystem.

Another important prediction of climate change is that these long periods of warmer weather will be interspersed with occasional major storms. These storms are predicted to be more intense both in magnitude and quantity (Korty 2022). Our results from immediately after Storm Carmel and the subsequent ‘stormy’ and cold winter of 2021-2022 represent a possible scenario for the effect of such major storms. In the winter of 2021 there was a major input of nutrients, particularly N+N (average ∼750 nM) and DIP (8.1 nM measured and a further 13.6 nM calculated to be present) into the photic zone. As a consequence, there was a substantial increase in the pico and nanophytoplankton biomass as well as strong circumstantial evidence of a major bloom of eukaryotes including diatoms. This autotrophic bloom created a much higher in-situ biomass than normal which represents a higher albeit temporary food source for the base of the food web; “Feast” conditions. Thus our study suggests that under conditions of anthropogenic climate change, these predicted changes in nutrient supply will become more extreme with longer periods of ‘Famine’ interspersed with occasional periods of ‘Feast’ caused by major winter storms.

## Acknowledgments

We thank the captains and crew of the R/V Mediterranean Explorer and R/V Bat Galim, the SoMMoS core sampling team and Eli Shemesh for help with the sampling. The ship-time was funded by the Leon H. Charney School of Marine Sciences with help from EcoOcean. The study is part of the Ph.D. work of Tal Ben Ezra. This study was conducted with support during data collection and analysis from the staff at the Morris Kahn Marine Research Station. This study was submitted under the umbrella of ocean@leeds group. We acknowledge support from the Helmholtz funded International Laboratory; The Eastern Mediterranean Sea Centre-An Early-Warning Model-System for our Future Oceans: EMS Future Ocean REsearch (EMS FORE).

## Sample Credit author statement

**Tal Ben-Ezra**: Conceptualization, Methodology, Sampling, Investigation, Writing –first draft, submission **Alon Blachinsky**: Methodology, Sampling, **Shiran Gozali**: Methodology Sampling, **Anat Tsemel**: Methodology Sampling, writing review and editing **Yotam Fadida:** Remote sensing, data interpretation, **Dan Tchernov**: Project administration, funding acquisition, conceptualisation, Student supervision **Yoav Lehahn**: Remote sensing, data interpretation, writing review and editing; **Tatiana Margo Tsagaraki**: XRF measurements and interpretation, writing review and editing; **Ilana Berman-Frank:** student supervision, writing review and editing; **Michael Krom:** Conceptualization, Sampling, Writing –first draft, student supervision.

## Supplementary material

**Table S1:** Sampling dates and parameters measured from research cruises between January 2020 and June 2022:

| Date | Frequency | Inorganic<br>nutrients<br>(N+N, DIP,<br>NH<sub>4</sub>, SiO<sub>2</sub>) | XRF | FCM | CTD | Notes / References |
| --- | --- | --- | --- | --- | --- | --- |
| February 2018<br>- January 2019 | Monthly | X |  | X | X | Ben-Ezra et al., 2021; Reich<br>et al., 2022 |
| January 2020 -<br>January 2021 | Monthly | X | X | X | X | XRF from Apr 2020 until Jan<br>2021 |
| February 2021<br>- June 2022 | Feb, Apr, Jun,<br>Aug, Dec 2021<br>Feb, Mar, Jun<br>2022 | X |  | X | X |  |

**Figure S1:**
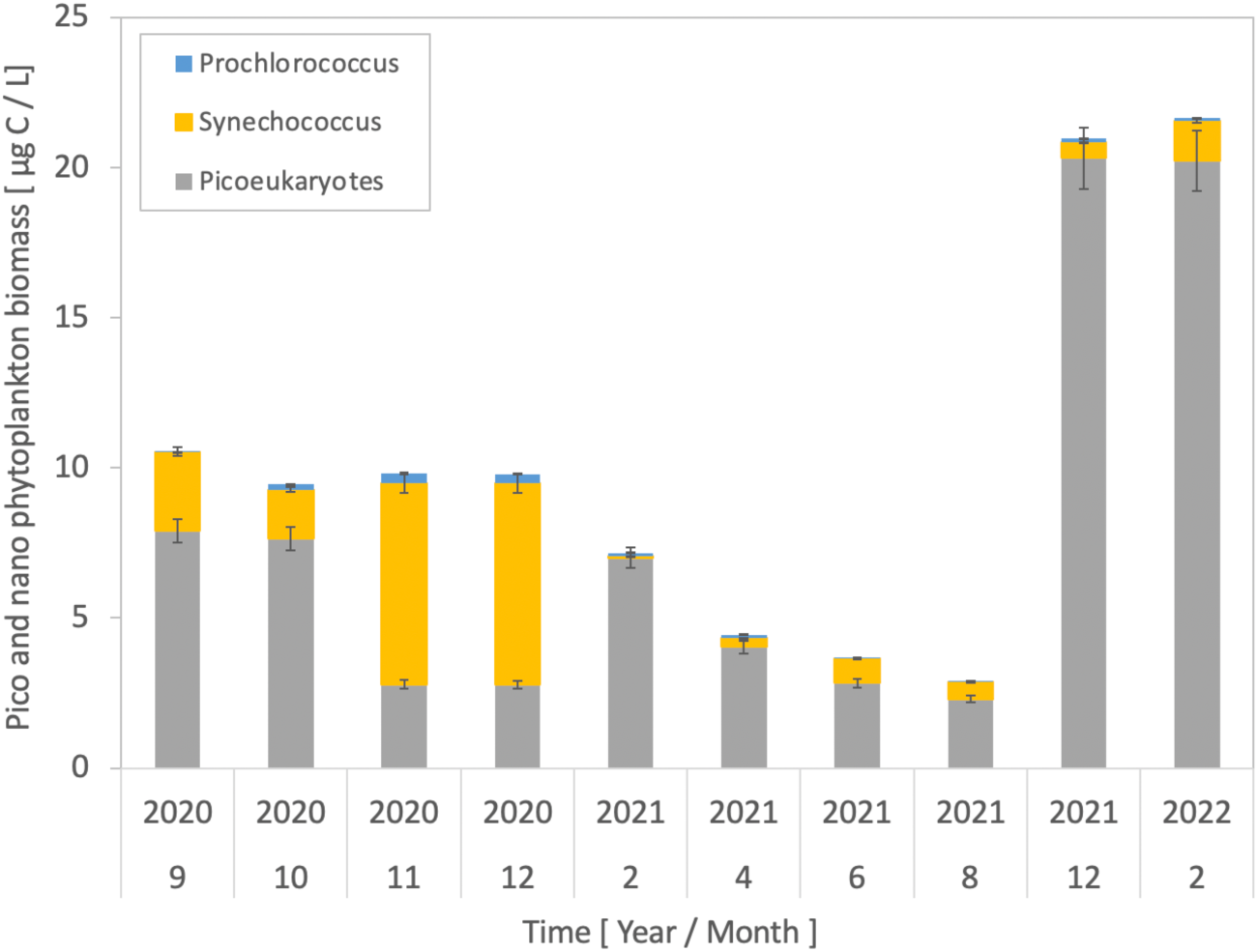
Calculated pico and nanophytoplankton biomass at 10 m depth. Temporal changes in the calculated biomass of Prochlorococcus, Synechococcus and picoeukaryotes at 10 m depth from September 2020 until February 2021, after Storm Carmel (17-24th of December 2021).

It was recognised that because were only able to determine a limited number of total water column biomass in late 2020 and through 2021 (Figure 8), it was appropriate to also present the samples taken in the water column at 10m from September 2020 to February 2022 (Figure 9). At 10 m depth the biomass of pico and nanoplankton during autumn 2020 was ∼10 □gC L^−1^ mainly as picoeukaryotes and *Synechococcus* with minimal *Prochlorococcus* biomass. In February 2021 the total pico and nanoplankton biomass had decreased to ∼7 □gC L^−1^ and continued to decrease through 2021 to a minimum of ∼2.8 □gC L^−1^ by August 2021, the last month sampled in 2021. Immediately after Storm Carmel, the total biomass increased by a factor of almost 10 to >20 □gC L^−1^ with picoeukaryotes now dominant. This is likely a conservative estimate, as it is probable that a significant biomass of larger eukaryotes was also present in the water column during this period (see discussion).

**Figure S2:**
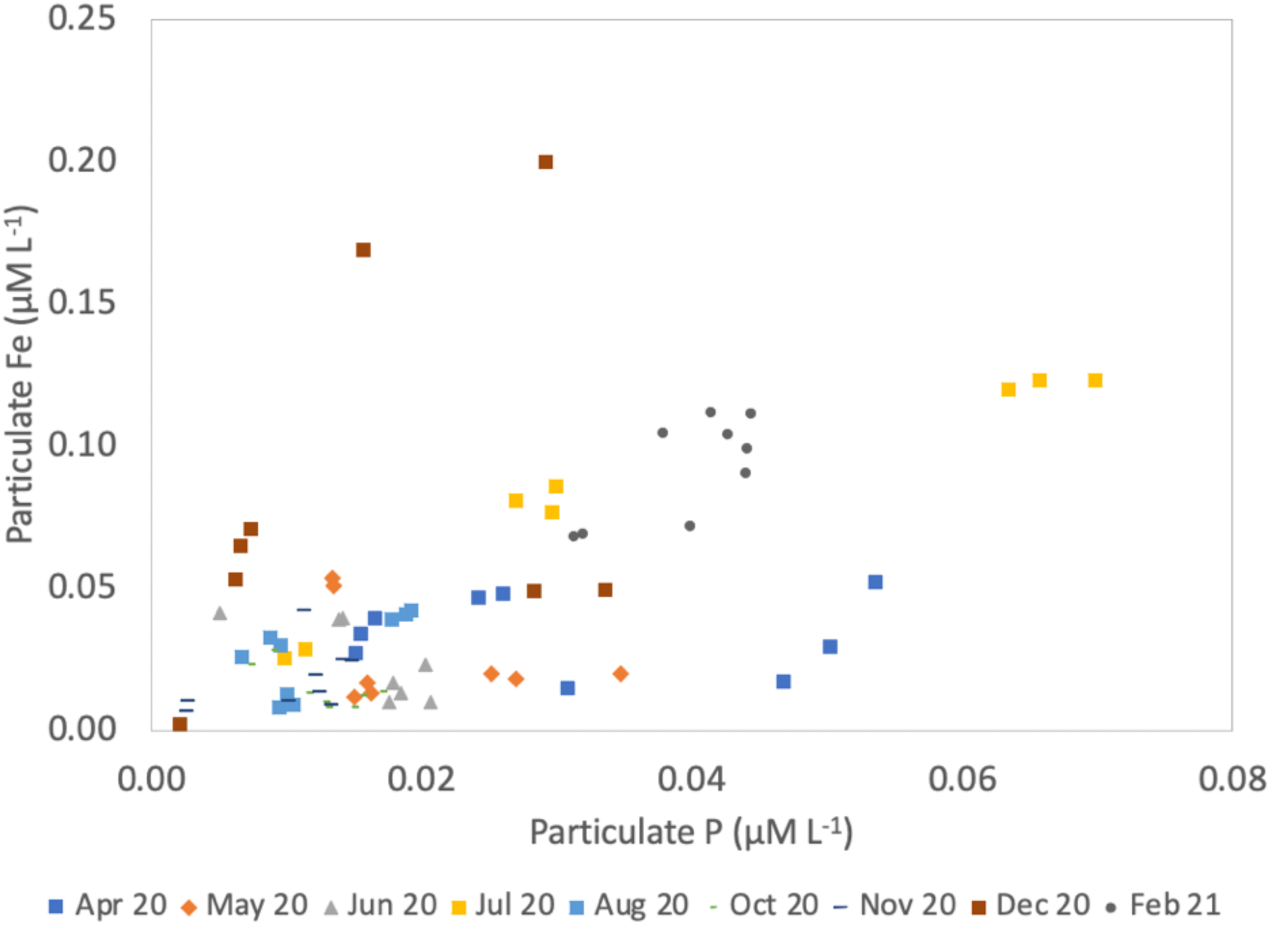
Particulate Fe vs Particulate P by WDXRF. Plot of Particulate Fe vs Particulate P (in μM L^−1^) from samples collected at 25, 100 and 180 m depth at N800 time-series station. The highest Fe: P ratio was 7.2:1 for December 2020 which had particles with unusually high Fe:Si ratio (Figure 5a). The Fe: P ratio was 1.54 for July 2020 when there was the maximum inorganic particles in the water column. At all other times the Fe: P ratio was similar (August and February) or lower i.e. there was an even greater proportion of P in the water column almost certainly as organic P.

**Table S2:** Missing DIP calculation. Measured concentration of DIP and N+N in the upper 200 965 m of the water column in August 2021 (during the period of seasonal stratification) compared with the measured concentration in December 2021 shortly after Storm Carmel. The values marked in red (December 2021 after Storm Carmel) and blue (August 2021) are the values for nutrients above the nutricline which were 100 m and 140 m, respectively.

| December 2021 shortly after Storm Carmel |  |  |  |  |
| --- | --- | --- | --- | --- |
| Depth (m) | Measured DIP<br>(nM) | Measured N+N<br>($\mu\text{M}$) | Calculated in-<br>tegrated DIP<br>($\mu\text{molesP m}^{-2}$) | Calculated inte-<br>grated N+N<br>($\text{mmolesN m}^{-2}$) |
| 2.5 | 2 | 1.09 | 4 | 2.17 |
| 10 | 6 | 0.73 | 48 | 5.88 |
| 25 | 1 | 0.76 | 15 | 11.46 |
| 50 | 3 | 0.61 | 75 | 15.18 |
| 75 | 2 | 0.67 | 50 | 16.68 |
| 100 | 10 | 0.69 | 250 | 17.35 |
| 125 | 9 | 0.68 | 225 | 17.15 |
| 150 | 11 | 0.68 | 275 | 16.95 |
| 160 | 8 | 0.82 | 80 | 8.21 |
| 180 | 12 | 1.38 | 240 | 27.68 |
| 200 | 116 | 5.58 | 2320 | 111.74 |
| Total amount of dissolved N+N or DIP in the photic zone |  |  | 817 | 96.09 |

| Depth (m) | Measured DIP<br>(nM) | Measured N+N<br>(μM) | Calculated in-<br>tegrated DIP<br>(μmolesP m<sup>-2</sup>) | Calculated inte-<br>grated N+N<br>(mmolesN m<sup>-2</sup>) |
| --- | --- | --- | --- | --- |
| 2.5 | 11 | 0.13 | 22 | 0.27 |
| 10 | 9 | 0.06 | 72 | 0.49 |
| 50 | 6 | 0.00 | 240 | 0.00 |
| 75 | 5 | 0.37 | 125 | 0.93 |
| 100 | 5 | 0.70 | 125 | 17.55 |
| 125 | 6 | 0.68 | 150 | 17.10 |
| 140 | 15 | 0.64 | 225 | 9.69 |
| 150 | 7 | 0.66 | 175 | 16.63 |
| 180 | 23 | 0.62 | 690 | 18.81 |
| 200 | 41 | 0.64 | 820 | 12.94 |
| Total amount of dissolved N+N or DIP in the photic zone |  |  | 459 | 1.68 |

## Reference List

1. Behrenfeld, M. J., R. T. O’Malley, D. A. Siegel, and others. 2006. Climate-driven trends in contemporary ocean productivity. Nature 444: 752–755.

2. Ben-Ezra, T., T. Reich, A. Tsemel, I. Berman-Frank, Y. Lehahn, D. Sher, Y. Suari, and M. D. Krom. 2023. Nutrient dynamics across the Israeli coastal shelf: An unusual oligotrophic coastal system. Cont. Shelf Res. 266: 105103. doi:10.1016/j.csr.2023.105103

3. Berthelot, H., S. Duhamel, S. L’Helguen, J. Maguer, and N. Cassar. 2021. Inorganic and organic carbon and nitrogen uptake strategies of picoplankton groups in the northwestern Atlantic Ocean. Limnol. Oceanogr. 66: 3682–3696.

4. Berthon, J.-F., and G. Zibordi. 2004. Bio-optical relationships for the northern Adriatic Sea. Int. J. Remote Sens. 25: 1527–1532.

5. Campbell, L. 2001. Flow cytometric analysis of autotrophic picoplankton. Methods Microbiol. 30: 317–343.

6. Carbo, P., M. D. Krom, W. B. Homoky, L. G. Benning, and B. Herut. 2005. Impact of atmospheric deposition on N and P geochemistry in the southeastern Levantine basin. Deep Sea Res. Part II Top. Stud. Oceanogr. 52: 3041–3053.

7. Casotti, R., A. Landolfi, C. Brunet, F. d’Ortenzio, O. Mangoni, M. Ribera d’Alcalà, and M. Denis. 2003. Composition and dynamics of the phytoplankton of the Ionian Sea (eastern Mediterranean). J. Geophys. Res. Ocean. 108.

8. Cavender-Bares, K. K., D. M. Karl, and S. W. Chisholm. 2001. Nutrient gradients in the western North Atlantic Ocean: Relationship to microbial community structure and comparison to patterns in the Pacific Ocean. Deep. Res. Part I Oceanogr. Res. Pap. 48: 2373–2395. doi:10.1016/S0967-0637(01)00027-9

9. Crombet, Y., K. Leblanc, B. Queguiner, and others. 2011. Deep silicon maxima in the stratified oligotrophic Mediterranean Sea.

10. D’Alimonte, D., F. Mélin, G. Zibordi, and J.-F. Berthon. 2003. Use of the novelty detection technique to identify the range of applicability of empirical ocean color algorithms. IEEE Trans. Geosci. Remote Sens. 41: 2833–2843.

11. D’Amario, B., C. Pérez, M. Grelaud, P. Pitta, E. Krasakopoulou, and P. Ziveri. 2020. Coccolithophore community response to ocean acidification and warming in the Eastern Mediterranean Sea: results from a mesocosm experiment. Sci. Rep. 10: 12637.

12. Denis, M., M. Thyssen, V. Martin, B. Manca, and F. Vidussi. 2010. Ultraphytoplankton basinscale distribution in the eastern Mediterranean Sea in winter: link to hydrodynamism and nutrients. Biogeosciences 7: 2227–2244.

13. Djaoudi, K., F. Van Wambeke, L. Coppola, and others. 2018a. Sensitive determination of the dissolved phosphate pool for an improved resolution of its vertical variability in the surface layer: New views in the P-depleted Mediterranean Sea. Front. Mar. Sci. 5: 1–11. doi:10.3389/fmars.2018.00234

14. Djaoudi, K., F. Van Wambeke, L. Coppola, and others. 2018b. Sensitive determination of the dissolved phosphate pool for an improved resolution of its vertical variability in the surface layer: New views in the P-depleted Mediterranean Sea. Front. Mar. Sci. 5. doi:10.3389/fmars.2018.00234

15. Eijsink, L. M., M. D. Krom, and B. Herut. 2000. Speciation and burial flux of phosphorus in the surface sediments of the eastern Mediterranean. Am. J. Sci. 300: 483–503.

16. Emeis, K., P. Mara, T. Schlarbaum, J. Möbius, K. Dähnke, U. Struck, N. Mihalopoulos, and M. Krom. 2010. External N inputs and internal N cycling traced by isotope ratios of nitrate, dissolved reduced nitrogen, and particulate nitrogen in the eastern Mediterranean Sea. J. Geophys. Res. Biogeosciences 115.

17. Ben Ezra, T., M. D. Krom, A. Tsemel, and others. 2021. Seasonal nutrient dynamics in the P depleted Eastern Mediterranean Sea. Deep. Res. Part I Oceanogr. Res. Pap. 176: 103607. doi:10.1016/j.dsr.2021.103607

18. Ganor, E., I. Osetinsky, A. Stupp, and P. Alpert. 2010. Increasing trend of African dust, over 49 years, in the eastern Mediterranean. J. Geophys. Res. Atmos. 115.

19. Guerzoni, S., R. Chester, F. Dulac, and others. 1999. The role of atmospheric deposition in the biogeochemistry of the Mediterranean Sea. Prog. Oceanogr. 44: 147–190.

20. Haber, M., D. Roth Rosenberg, M. Lalzar, and others. 2022. Spatiotemporal variation of microbial communities in the ultra-oligotrophic Eastern Mediterranean Sea. Front. Microbiol. 13: 867694.

21. Hurd, C. L., A. Lenton, B. Tilbrook, and P. W. Boyd. 2018. Current understanding and challenges for oceans in a higher-CO2 world. Nat. Clim. Chang. 8: 686–694.

22. Ignatiades, L., S. Psarra, V. Zervakis, K. Pagou, E. Souvermezoglou, G. Assimakopoulou, and O. Gotsis-Skretas. 2002. Phytoplankton size-based dynamics in the Aegean Sea (Eastern Mediterranean). J. Mar. Syst. 36: 11–28. doi:10.1016/S0924-7963(02)00132-X

23. Kamennaya, N. A., K. Geraki, D. J. Scanlan, and M. V. Zubkov. 2020. Accumulation of ambient phosphate into the periplasm of marine bacteria is proton motive force dependent. Nat. Commun. 11: 1–13. doi:10.1038/s41467-020-16428-w

24. Keuter, S., J. Silverman, M. D. Krom, and others. 2022. Seasonal patterns of coccolithophores in the ultra-oligotrophic South-East Levantine Basin, Eastern Mediterranean Sea. Mar. Micropaleontol. 102153. doi:10.1016/J.MARMICRO.2022.102153

25. Korty, R. L. 2022. Seas reveal a surge in the strength of tropical storms.

26. Krause, J. W., M. W. Lomas, and D. M. Nelson. 2009. Biogenic silica at the Bermuda Atlantic Time-series Study site in the Sargasso Sea: Temporal changes and their inferred controls based on a 15-year record. Global Biogeochem. Cycles 23.

27. Kress, N., and B. Herut. 2001. Spatial and seasonal evolution of dissolved oxygen and nutrients in the Southern Levantine Basin (Eastern Mediterranean Sea): chemical characterization of the water masses and inferences on the N: P ratios. Deep Sea Res. Part I Oceanogr. Res. Pap. 48: 2347–2372.

28. Krom, M. D., S. Brenner, N. Kress, A. Neori, and L. I. Gordon. 1992. Nutrient dynamics and new production in a warm-core eddy from the Eastern Mediterranean Sea. Deep Sea Res. Part A. Oceanogr. Res. Pap. 39: 467–480.

29. Krom, M. D., K. C. Emeis, and P. Van Cappellen. 2010. Why is the Eastern Mediterranean phosphorus limited? Prog. Oceanogr. 85: 236–244. doi:10.1016/j.pocean.2010.03.003

30. Krom, M. D., N. Kress, I. Berman-Frank, and E. Rahav. 2014a. Past, present and future patterns in the nutrient chemistry of the eastern mediterranean, p. 49–68. *In* The Mediterranean Sea: Its History and Present Challenges. Springer Netherlands.

31. Krom, M. D., N. Kress, S. Brenner, and L. I. Gordon. 1991. Phosphorus limitation of primary productivity in the eastern Mediterranean Sea. Limnol. Oceanogr. 36: 424–432. doi:10.4319/lo.1991.36.3.0424

32. Krom, M. D., N. Kress, and K. Fanning. 2014b. Silica cycling in the ultra-oligotrophic eastern Mediterranean Sea. Biogeosciences 11: 4211–4223.

33. Krom, M. D., Z. Shi, A. Stockdale, and others. 2016. Response of the Eastern Mediterranean microbial ecosystem to dust and dust affected by acid processing in the atmosphere. Front. Mar. Sci. 3: 133.

34. Krom, M. D., T. F. Thingstad, S. Brenner, and others. 2005a. Summary and overview of the CYCLOPS P addition Lagrangian experiment in the Eastern Mediterranean. Deep. Res. Part II Top. Stud. Oceanogr. 52: 3090–3108. doi:10.1016/j.dsr2.2005.08.018

35. Krom, M. D., E. M. S. Woodward, B. Herut, and others. 2005b. Nutrient cycling in the south east Levantine basin of the eastern Mediterranean: Results from a phosphorus starved system. Deep. Res. Part II Top. Stud. Oceanogr. 52: 2879–2896. doi:10.1016/j.dsr2.2005.08.009

36. Lomas, M. W., N. R. Bates, R. J. Johnson, A. H. Knap, D. K. Steinberg, and C. A. Carlson. 2013. Two decades and counting: 24-years of sustained open ocean biogeochemical measurements in the Sargasso Sea. Deep Sea Res. Part II Top. Stud. Oceanogr. 93: 16–32. doi:10.1016/J.DSR2.2013.01.008

37. Marinov, I., S. C. Doney, and I. D. Lima. 2010. Response of ocean phytoplankton community structure to climate change over the 21st century: partitioning the effects of nutrients, temperature and light. Biogeosciences 7: 3941–3959.

38. Masuda, T., K. Inomura, J. Mareš, and others. 2023. Coexistence of dominant marine phytoplankton sustained by nutrient specialization. Microbiol. Spectr. 11: e04000–22.

39. Moutin, T., and P. Raimbault. 2002. Primary production, carbon export and nutrients availability in western and eastern Mediterranean Sea in early summer 1996 (MINOS cruise). J. Mar. Syst. 33: 273–288.

40. Moutin, T., F. Van Wambeke, and L. Prieur. 2012. Introduction to the Biogeochemistry from the Oligotrophic to the Ultraoligotrophic Mediterranean (BOUM) experiment. Biogeosciences 9: 3817–3825.

41. Mulas, M., J. Silverman, T. Guy-Haim, S. Noe, and G. Rilov. 2022. Thermal vulnerability of the Levantine endemic and endangered habitat-forming macroalga, Gongolaria rayssiae: implications for reef carbon. Front. Mar. Sci. 9: 862332.

42. Murphy, J., and J. P. Riley. 1962. A modified single solution method for the determination of phosphate in natural waters. Anal. Chim. Acta 27: 31–36. doi:10.1016/S0003-2670(00)88444-5

43. Ozer, T., I. Gertman, H. Gildor, and B. Herut. 2022. Thermohaline Temporal Variability of the SE Mediterranean Coastal Waters (Israel) – Long-Term Trends, Seasonality, and Connectivity. Front. Mar. Sci. 8: 1–14. doi:10.3389/fmars.2021.799457

44. Ozer, T., I. Gertman, N. Kress, J. Silverman, and B. Herut. 2017. Interannual thermohaline (1979–2014) and nutrient (2002–2014) dynamics in the Levantine surface and intermediate water masses, SE Mediterranean Sea. Glob. Planet. Change 151: 60–67.

45. Paulino, A. I., M. Heldal, S. Norland, and J. K. Egge. 2013. Elemental stoichiometry of marine particulate matter measured by wavelength dispersive X-ray fluorescence (WDXRF) spectroscopy. J. Mar. Biol. Assoc. United Kingdom 93: 2003–2014.

46. Powley, H. R., M. D. Krom, and P. Van Cappellen. 2017. Understanding the unique biogeochemistry of the Mediterranean Sea: Insights from a coupled phosphorus and nitrogen model. Global Biogeochem. Cycles 31: 1010–1031. doi:10.1002/2017GB005648

47. Powley, H. R., M. D. Krom, K.-C. Emeis, and P. Van Cappellen. 2014. A biogeochemical model for phosphorus and nitrogen cycling in the Eastern Mediterranean Sea: Part 2. Response of nutrient cycles and primary production to anthropogenic forcing: 1950–2000. J. Mar. Syst. 139: 420–432.

48. Psarra, S., T. Zohary, M. D. Krom, and others. 2005. Phytoplankton response to a Lagrangian phosphate addition in the Levantine Sea (Eastern Mediterranean). Deep. Res. Part II Top. Stud. Oceanogr. 52: 2944–2960. doi:10.1016/j.dsr2.2005.08.015

49. Pujo-Pay, M., P. Conan, L. Oriol, and others. 2011. Integrated survey of elemental stoichiometry (C, N, P) from the western to eastern Mediterranean Sea. Biogeosciences 8: 883–899. doi:10.5194/bg-8-883-2011

50. Reich, T., T. Ben-Ezra, N. Belkin, and others. 2022. A year in the life of the Eastern Mediterranean: Monthly dynamics of phytoplankton and bacterioplankton in an ultraoligotrophic sea. Deep. Res. Part I Oceanogr. Res. Pap. 182: 103720. doi:10.1016/j.dsr.2022.103720

51. Reynolds, S., C. Mahaffey, V. Roussenov, and R. G. Williams. 2014. Evidence for production and lateral transport of dissolved organic phosphorus in the eastern subtropical North Atlantic. Global Biogeochem. Cycles 28: 805–824.

52. Roether, W., B. Klein, V. Beitzel, and B. B. Manca. 1998. Property distributions and transient-tracer ages in Levantine Intermediate Water in the Eastern Mediterranean. J. Mar. Syst. 18: 71–87.

53. Schlitzer, and Reiner. 2023. Ocean Data View.

54. Schroeder, K., J. Chiggiato, S. A. Josey, M. Borghini, S. Aracri, and S. Sparnocchia. 2017. Rapid response to climate change in a marginal sea. Sci. Rep. 7: 4065.

55. SEAL Analytical. 2011a. Ammonia in water and seawater. AutoAnalyzer Appl. G-327–05.

56. SEAL Analytical. 2011b. Nitrate and Nitrite in water and seawater Total Nitrogen in persulfate digests. AutoAnalyzer Appl. G-172–96.

57. SEAL Analytical. 2011c. Silicate in Water and Seawater Method-No. G-177-96. AutoAnalyzer Appl. G-177–96.

58. Shan, K., Y. Lin, P.-S. Chu, X. Yu, and F. Song. 2023. Seasonal advance of intense tropical cyclones in a warming climate. Nature 1–7.

59. Steinberg, D. K., C. A. Carlson, N. R. Bates, R. J. Johnson, A. F. Michaels, and A. H. Knap. 2001. Overview of the US JGOFS Bermuda Atlantic Time-series Study (BATS): a decade-scale look at ocean biology and biogeochemistry. Deep Sea Res. Part II Top. Stud. Oceanogr. 48: 1405–1447.

60. Stockdale, A., M. D. Krom, R. J. G. Mortimer, and others. 2016. Understanding the nature of atmospheric acid processing of mineral dusts in supplying bioavailable phosphorus to the oceans. Proc. Natl. Acad. Sci. 113: 14639–14644.

61. Theodosi, C., Z. Markaki, F. Pantazoglou, A. Tselepides, and N. Mihalopoulos. 2019. Chemical composition of downward fluxes in the Cretan Sea (Eastern Mediterranean) and possible link to atmospheric deposition: A 7 year survey. Deep Sea Res. Part II Top. Stud. Oceanogr. 164: 89–99.

62. Thingstad, T. F., M. D. Krom, R. F. C. Mantoura, and others. 2005. Nature of phosphorus limitation in the ultraoligotrophic eastern Mediterranean. Science (80-.). 309: 1068–1071. doi:10.1126/science.1112632

63. Vichi, M., W. May, and A. Navarra. 2003. Response of a complex ecosystem model of the northern Adriatic Sea to a regional climate change scenario. Clim. Res. 24: 141–159.

64. Vidussi, F., H. Claustre, B. B. Manca, A. Luchetta, and J. Marty. 2001. Phytoplankton pigment distribution in relation to upper thermocline circulation in the eastern Mediterranean Sea during winter. J. Geophys. Res. Ocean. 106: 19939–19956.

65. Volpe, G., S. Colella, V. E. Brando, and others. 2019. Mediterranean ocean colour Level 3 operational multi-sensor processing. Ocean Sci. 15: 127–146.

66. Volpe, G., B. B. Nardelli, S. Colella, A. Pisano, and R. Santoleri. 2018. Operational Interpolated Ocean Colour Product in the Mediterranean Sea. New Front. Oper. Oceanogr. 227–244.

67. Winder, M., and U. Sommer. 2012. Phytoplankton response to a changing climate. Hydrobiologia 698: 5–16.

68. Wu, J., W. Sunda, E. A. Boyle, and D. M. Karl. 2000. Phosphate depletion in the Western North Atlantic Ocean. Science (80-.). 289: 759–762. doi:10.1126/science.289.5480.759

69. Zohary, T., B. Herut, M. D. Krom, and others. 2005. P-limited bacteria but N and P co-limited phytoplankton in the Eastern Mediterranean – A microcosm experiment. Deep. Res. Part II Top. Stud. Oceanogr. 52: 3011–3023. doi:10.1016/j.dsr2.2005.08.011

70. Zohary, T., and R. R. D. Robarts. 1998. Experimental study of microbial P limitation in the eastern Mediterranean. Limnol. Oceanogr. 43: 387–395. doi:10.4319/lo.1998.43.3.0387

71. Zubkov, M. V, A. P. Martin, M. Hartmann, C. Grob, and D. J. Scanlan. 2015. Dominant oceanic bacteria secure phosphate using a large extracellular buffer. Nat. Commun. 6: 7878.

72. Zubkov, M. V, I. Mary, E. M. S. Woodward, P. E. Warwick, B. M. Fuchs, D. J. Scanlan, and P. H. Burkill. 2007. Microbial control of phosphate in the nutrient-depleted North Atlantic subtropical gyre. Environ. Microbiol. 9: 2079–2089.

73. Windguru Station 2049.

